# Energy landscapes of A*β* monomers are sculpted in accordance with Ostwald’s rule of stages

**DOI:** 10.1101/2022.06.16.496424

**Authors:** Debayan Chakraborty, John E. Straub, D Thirumalai

**Affiliations:** Department of Chemistry, The University of Texas at Austin, 24th Street Stop A5300, Austin TX 78712, USA; Department of Chemistry, Boston University, Massachusetts 022155, USA

## Abstract

The transition from a disordered to an assembly-competent and sparsely populated monomeric state (N*) in amyloidogenic sequences is a crucial event in the aggregation cascade. Using a well-calibrated model for Intrinsically Disordered Proteins (IDPs), we show that the N* states, which bear considerable resemblance to distinct polymorphic fibril structures found in experiments, not only appear as excitations on the monomer free energy landscapes of A*β*40 and A*β*42 but also initiate the aggregation cascade. Interestingly, for *A*β**42, the transitions to the different N* states are in accord with Ostwald’s rule of stages, with the least stable structures forming ahead of thermodynamically favored structures, which appear only on longer time-scales. Despite having similar topographies, the A*β*40 and A*β*42 monomer landscapes exhibit different extent of ruggedness, particularly in the vicinity of N* states, which we show have profound implications in dictating the intramolecular diffusion rates, and subsequent self-assembly into higher order structures. The network of connected kinetic states, which for A*β*42 is considerably more complex than for A*β*40, shows that the most favored dimerization routes proceed via the N* states. Direct transition between the disordered ground states within the monomer and dimer basins is less likely. The Ostwald’s rule of stages holds widely, qualitatively explaining the unusual features in other fibril forming IDPs, such as Fused in Sarcoma (FUS). Similarly, the N* theory accounts for dimer formation in small disordered polyglutamine peptides, implicated in the Huntington disease.

**Graphical TOC Entry:** 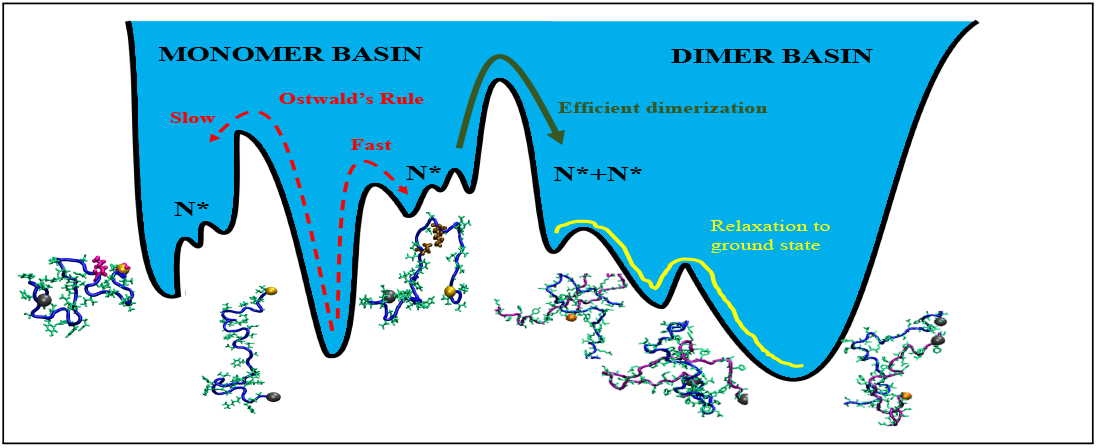

## Introduction

The accumulation of insoluble plaques and neurofibrillary tangles inside different parts of the central nervous system is a hallmark of Alzheimer’s disease (AD), one of the well-known proteinopathies.^1–5^ The amyloid plaques have a characteristic cross-β architecture and are primarily composed of A*β* peptides of length 39-43 residues.^6^ These peptides are produced *in vivo* by the proteolytic cleavage of the nearly 770-residue amyloid precursor protein (APP) by γ-secretases,^7,8^ with A*β*40 and A*β*42 being the most common cleavage products. Despite having globally similar topologies, A*β* fibrils exhibit diverse structural polymorphism.^9–11^ The specific structure that is populated could depend on the differences in the fibril growth conditions,^12^ as well as preferential hydration of certain polymorphs.^13^ Although A*β* fibrils were considered to be the key players in AD etiology,^14,15^ recent works suggest that the oligomers^16–18^ formed early along the aggregation cascade could be the real culprits. Other neurodegenerative disorders, such as Parkinson’s disease (PD) and Huntington’s disease, which also result from aberrant protein aggregation, may share certain common themes with AD,^19,20^ including mechanisms of fibril formation, polymorphic fibril morphologies, as well the association of cytotoxicity with oligomers. In order to uncover the general principles, a microscopic understanding of the key steps connecting the monomer to the fibril state, is necessary. ^21,22^

The first step in the aggregation cascade involves the conformational transitions of the A*β* monomer to an assembly-competent structure (henceforth, referred to as the N* state), having some structural signatures of the fibril state. The N* states are excitations on the monomer free energy landscape.^23–29^ Because the N* states are only sparsely populated (typically less than 5%) under normal growth conditions, they can only be resolved by experiments with high spatial and temporal resolution. ^30,31^ In this context, computer simulations are useful in directly detecting these “dark states”. The different N* states could self-assemble to form oligomers of different sizes. A critical nucleus once formed, serves as a template for protofibril formation, and it continues to grow through the deposition of monomers via a ‘dock-lock’ mechanism.^21^ In this model of protein aggregation,^21,24^ the N* concept plays a central role. It not only accounts for the possibility of fibril polymorphism, but also provides a basis for understanding the aggregation propensities of different peptide sequences.^29,32^

The A*β*40 and A*β*42 sequences, which are the major isoforms implicated in AD, behave like intrinsically disordered proteins in their monomeric forms, ^31,33,34^ which is manifested by a lack of persistent secondary structures. Hence, a microscopic characterization of their conformational ensembles is a challenging task. Toward this end, computer simulations^35–39^ have provided important insights into structural ensembles of A*β* monomers, although the precise details and the corresponding ensemble-averages strongly depend on the force-field and the sampling strategy. Using the self-organized polymer model for intrinsically disordered proteins (SOP-IDP), we recently showed that the ensemble-averages cannot be used to distinguish between the aggregation propensities of A*β*40 and A*β*42 because much of the statistical weight is dominated by the random-coil like structures.^29^ In this sense, A*β*40 and A*β*42 behave like Flory random coil, with *R_g_ ~ a_0_N^ν^* (v ≈ 0.6), where *N* is the number of amino acids. It is only when the population of the different fibril-like states were identified using clustering techniques and geometric order parameters, the nearly one order of magnitude difference between the aggregation rates of A*β*40 and A*β*42 could be rationalized. Therefore, our work,^29^ underscores the importance of quantitatively characterizing the N* states present within the monomer conformational ensemble of assembly-competent IDPs in order to provide a microscopic basis for protein aggregation.^29,32^

Here, we construct the free energy landscapes of A*β*40 and A*β*42 using simulations of the SOP-IDP model. The landscapes exhibit a pronounced bias towards the disordered random coil ground state, rationalizing why much of the thermodynamic ensemble-averages are reminiscent of random-coils. A diverse range of N* states are encoded as excitations on the energy landscape, and they exhibit the key structural features found in different A*β*40 and A*β*42 polymorphs, such as the U-bend^40^ and the S-bend motifs.^41^ Despite having similar topographies, the landscapes are associated with different extent of ruggedness, particularly in regions where the probability to find the N* structures is the maximum. This finding has important implications in the intramolecular diffusivity of A*β* monomers, and subsequently their ability to coalesce with other binding partners. The dynamics of A*β* monomers is hierarchically organized, with relaxation time-scales in the sub-*μ*s regime, in agreement with recent experiments. ^42,43^ The transitions between the disordered configurations and the N* states are faster in A*β*42, suggesting that it is kinetically more predisposed to aggregate compared to A*β*40. Although there are multiple pathways to dimer formation, the most productive route is the one when both the monomers are in the N* state. Interestingly, the order of transitions to the different N* states in A*β*42 is in accord with Ostwald’s rule of stages, with the thermodynamically less favored U-bend structure appearing before the S-bend conformation. Thus, thermodynamic stability and fibril formation rates are inversely correlated, which we argue also prevails in the formation of fibrils in Fused in Sarcoma (FUS), and the associated variants and in polyglutamine peptides.

## Results

### Free energy landscapes as transition disconnectivity graphs (TRDGs)

The conformational spaces of the A*β* monomers were partitioned into distinct clusters (free energy minima) based on the distribution of reciprocal interatomic distances (DRID) metric. As described in earlier work,^44^ the DRID-based metric preserves the kinetic distances among different minima. The optimal number of clusters for A*β*40 and A*β*42 were identified using a knee-point analysis (see Supporting Information, and Fig. S3). The effective free energy barriers between different minima were estimated using the min-cut procedure, ^45^ which is based on the Ford and Fulkerson theorem, ^46^ and exploits the isomorphism between a network representation of the conformational landscape and a graph with capacitated edges (see Supporting Information for further details).

The free energy landscapes of A*β* monomers at 298 K are depicted in the form of TRDGs^45,47,48^ in Fig. 1 and Fig. 2. In contrast to other formulations that rely on low-dimensional projections onto pre-defined order parameters, the TRDG representation is a faithful representation of the underlying kinetics. In this “tree” structure^49^ (or equivalently the “kinetic transition network” representation),^50,51^ the landscape is partitioned into disjoint free energy basins, such that minima within each basin are mutually accessible, whereas inter-basin transitions only occur over longer observation time-scales (see Supporting Information for further details). In our case, each node (free energy minimum) in the TRDG (Fig. 1 and Fig. 2) corresponds to an ensemble of structures that undergo rapid interconversion through local fluctuations. The TRDG, therefore, is an effective representation of the structural heterogeneity of A*β* monomers, ^29^ as well as the hierarchical organization of their conformational dynamics. ^43^ Both features are known to mediate the early events along the aggregation cascade. The branches of the TRDGs (Figs. 1 and 2) are color-coded from red to blue to reflect the degree of structural similarity with respect to the experimental fibril structures (see captions in Fig. 1 and Fig. 2 for details).

**Figure 1.**
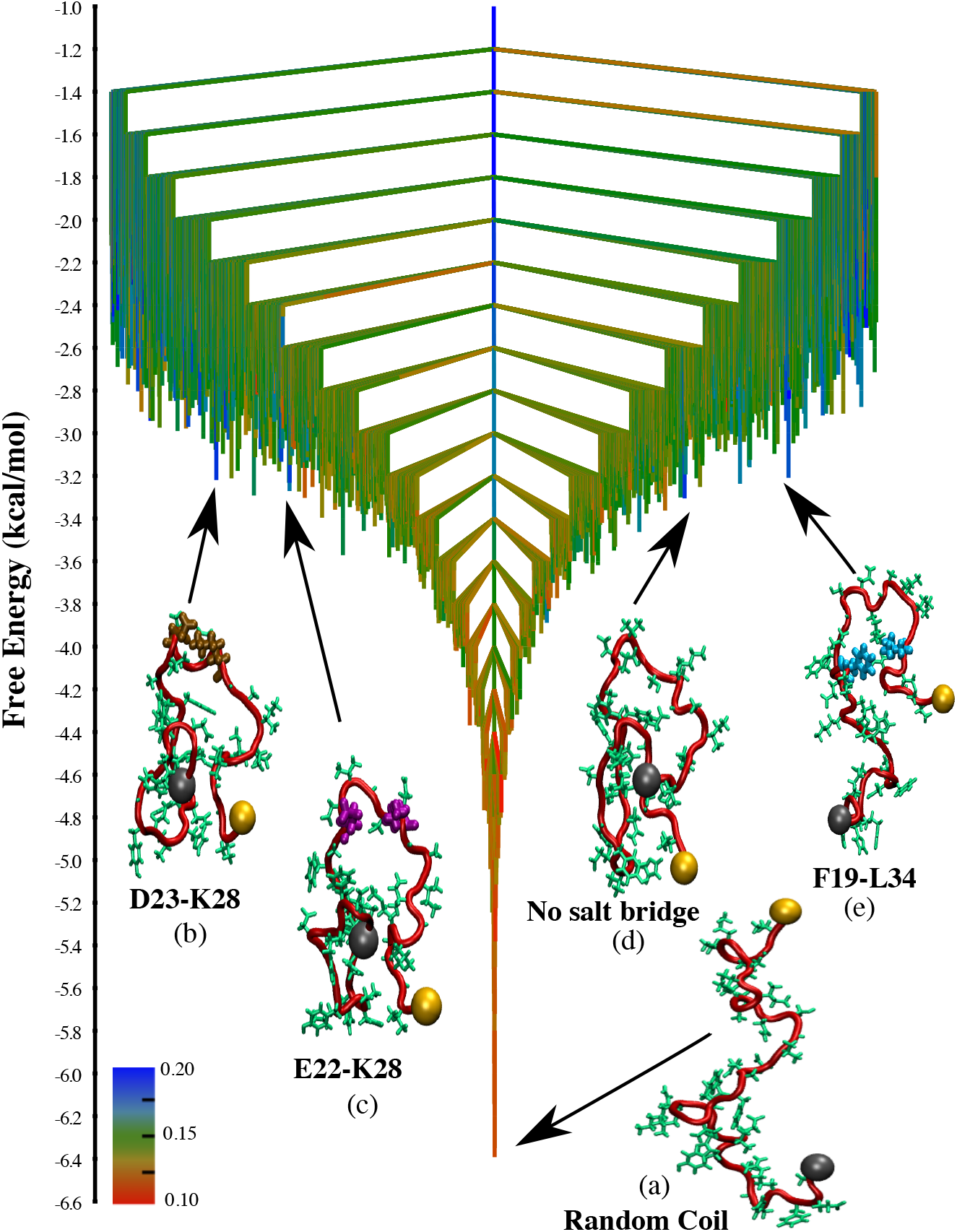
The free energy landscape for the A*β*40 monomer at 298 K depicted in the form of a transition disconnectivity graph (TRDG). The branches are colored according to 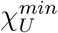, which is the minimum value of the overlap with respect to the experimental U-bend fibril structure, for a group of conformations constituting a free energy minimum (the scales quantifies the structural similarity scale). The N-terminus of the peptide is shown as a grey sphere, and the C-terminus is shown as an orange sphere. The key contacts within these structures are shown in different colors: D23-K28 salt-bridge (ochre); E22-K28 salt-bridge (purple); F19-L34 contact (cyan). An ensemble of random-coil (RC) structures corresponds to the free energy global minimum (a). The landscape exhibits minimal frustration (‘palmtree’ like features), suggesting that relaxation to the global minimum from other regions of the landscape occur very efficiently. Some representative snapshots corresponding to the different N* states (free energy excitations that resemble the fibril structure) are also shown. (b) A strand-loop-strand (SLS) topology stabilized by a VGSN turn, and a D23-K28 saltbridge. (c) SLS structure stabilized by a VGSN turn, and a E22-K28 salt-bridge. (d) SLS structure lacking the E22/D23-K28 salt-bridge. (e) SLS structure with a contact between residues F19 and L34.

**Figure 2.**
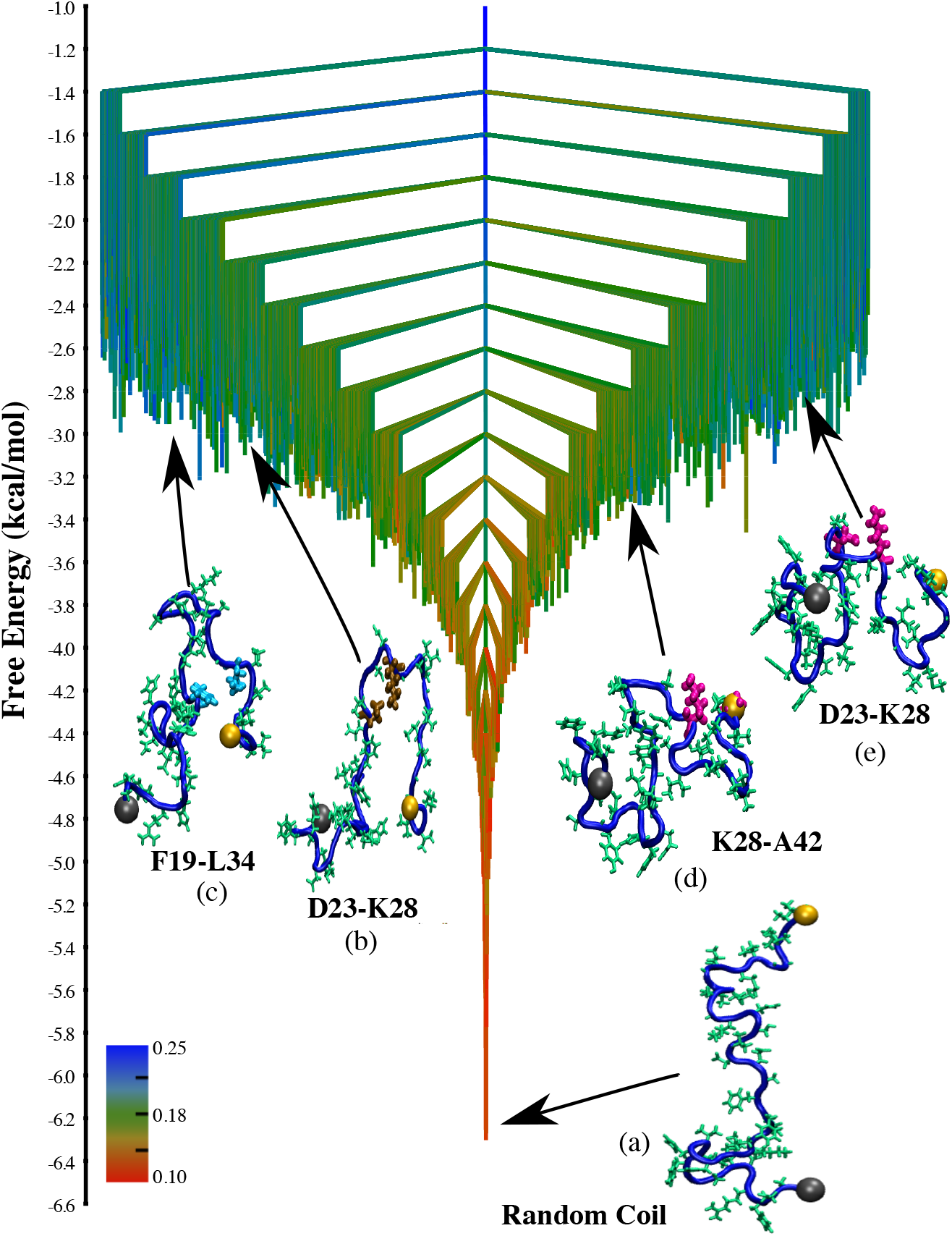
The free energy landscape for the A*β*42 monomer computed at 298 K depicted in the form of a transition disconnectivity graph (TRDG). The branches of the TRDG are color-coded according to either 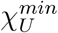 or 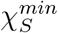, whichever is greater for a given free energy minimum. Here, 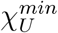 is the minimum value of the overlap with respect to the U-bend fibril, and 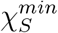 is the minimum value of the overlap with respect to the S-bend fibril, for conformations within a free energy minimum. The N-terminus of the peptide is shown as a grey sphere, and the C-terminus is shown as an orange sphere. Random-coil structures (RC) constitute the free energy global minimum (a), and the landscape appears minimally frustrated (similar to A*β*40). Some representative snapshots corresponding to the different *N*\* states are also shown. (b) SLS-like structure consisting of a U-bend near the VGSN turn region, and stabilized by a D23-K28 salt-bridge (shown in ochre). (c) SLS structure, stabilized by a F19-L34 contact (shown in cyan). (d) S-bend motif stabilized by a K28-A42 contact (shown in magenta). (e) S-bend motif having the K28-A42 contact replaced by a D23-K28 salt-bridge.

For both A*β*40 and A*β*42, random-coils (RC) devoid of any persistent structures represent the free energy ground state, and appear at the bottom of the energy landscapes (Fig. 1 and Fig. 2). The branches near the bottom of the TRDGs are colored red, indicating that conformations within the lowest free energy basins do not exhibit any structural similarity with respect to the monomers within the fibril states. For both A*β*40 and A*β*42, random-coils (RC) devoid of any persistent structures represent the free energy ground state, and appear at the bottom of the energy landscapes (Fig. 1 and Fig. 2). Hence, RCs would largely determine the thermodynamic properties (in other words, the experimentally measurable observables) at low temperatures. It is therefore not surprising that previous studies have predicted nearly negligible secondary structure propensities for A*β*40 and A*β*42 peptides,^29,33^ as well as practically identical ensemble averages for global chain dimensions. ^29,42^

The color of branches continuously changes from red to blue, with uphill excursions on the TRDGs. In other words, the essential signatures of the fibril-like order encoded within the free energy excitations (*N*\* states) of the monomer ensembles of A*β*40 and A*β*42 become apparent, despite the evident bias towards RC-like conformations at equilibrium. Some representative free energy minima, which exhibit the key structural elements of the fibril state, are shown superimposed on the TRDGs (Fig. 1 and Fig. 2). These conformations are only sparsely populated relative to the disordered state (with populations ≈ 5% or less),^26,29,30^ and their structural details are described below.

#### A*β*40

Several previous studies have suggested that the early events in the amyloid aggregation cascade critically depend on the presence of a D23-K28 salt-bridge,^23,52–54^ which stabilizes the VGSN turn region. Conformations within free energy minimum (b) (Fig. 1) clearly exhibit these contacts, and form the strand-loop-strand (SLS) structure, which forms the repeating unit in the U-bend A*β*40 fibril.^52^ Structures within minimum (c) (Figure 1) also exhibit the same SLS topology, but consist of a salt-bridge between residues E22 and K28. Statistical analysis from our earlier study showed that the D23-K28 salt-bridge is only marginally favored compared to E22-K28.^29^ Here, we find that the corresponding free energy minima are approximately isoenergetic. In addition to (*b*) and (*c*), where nearly all the constituent structures have the E22/D23-K28 salt-bridges we also find minima, which are structurally more heterogeneous. For instance, in the minimum (*d*) (Figure 1) all the structures exhibit the SLS topology, but the E22/D23-K28 salt-bridges are absent in a majority of them. The appearance of such structures on the free energy landscape hints at a scenario where the A*β*40 monomer could adopt the fibril-like topologies at the monomer level, with the D23-K28 contact appearing late during the aggregation cascade. This possibility was raised recently,^55^ where using solid-state NMR spectroscopy the authors speculated that the D23-K28 salt-bridge does not form at the monomer or the early oligomer stage.

In addition to the E22/D23-K28 salt-bridges, we also find many SLS topologies exhibiting a contact between the F19 and L34 residues (Fig. 1, snapshot (e)). Previous studies have shown that the perturbations of the F19-L34 can completely abrogate the cytotoxicity of A*β*40 fibrils,^56,57^ without inducing significant distortions in the fibril structure. Interestingly, experiments based on solid-state NMR spectroscopy^56–58^ indicate that unlike the D23-K28 salt-bridge which may be absent in early assembly intermediates, the F19-L34 contact persists throughout the aggregation cascade.

#### A*β*42

Similar to A*β*40, we also find a diverse range of excited states on the free energy landscape of A*β*42, exhibiting fibril-like morphologies. Free energy minimum (*b*) (Fig. 2) consists of structures having the canonical U-bend (SLS topology), and a salt-bridge between residues D23 and K28. These structures resemble the repeating units found in some A*β*42 polymorphs.^59^ Just as in A*β*40, we also identify free energy minima in the intermediate sections of the TRDG having a SLS topology stabilized by a F19-L34 contact (Fig. 2, snapshot (c)).

Besides the U-bend polymorph, A*β*42 also forms fibril structures in which the building block is a S-bend motif. Recent NMR^41,60^ and cryo-EM^61^ experiments have shown the S-bend results from enhanced structural ordering near the C-terminus of A*β*42, and is characterized by a hydrophobic contact between residues K28 and A42. Exclusive formation of the S-bend motif in case of A*β*42 could be linked to its higher aggregation propensity as compared to A*β*40.^29,62^ All the structures within the free energy minimum (*d* in Fig. 2) have the characteristic features of the S-bend motif, and exhibit a stable contact between residues K28 and A42. The S-bend structures within *e* (Fig. 2) lack the K28-A42 contact, and are substantially destabilized, suggesting that hydrophobic interactions between the terminal residues are critical for maintaining the complex topology. In agreement with our recent study,^29^ where we estimated the relative populations of different A*β*42 conformational ensembles based on structural clustering, we find that the S-bend topology having the K28-A42 contact has a lower free energy compared to the U-bend structures.

#### Relaxation time-scales

We estimated the time-scales associated with the relaxation of the different states, corresponding to the nodes in the TRDGs, to their equilibrium distributions. We assume that the kinetics may be described in terms of a discrete time Markov chain. ^63–65^ For a given vector, **P**(**t**), whose elements are the probabilities of finding the system in different states at time *t*, the following holds for Markovian dynamics:

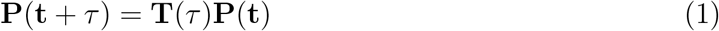

In Eq. (1) **T**(*τ*) is the transition matrix, whose elements *T_ij_*s are the probabilities of finding the system in state *j* at time *τ*, when it was in state *i* at time *t*. The highest eigenvalue of *T*(*τ*) is exactly 1, and the corresponding eigenvector represents the populations of the different states at thermal equilibrium. The other eigenvalues, λ_*i*_ are related to the implied relaxation time-scales, *t_i_* in the following way:

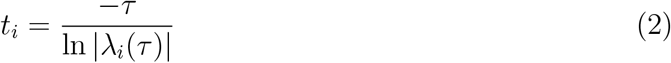

We find that the for both A*β*40, and A*β*42, the implied time-scales, *t_i_* are largely independent of the lag time, *τ* (see Supporting Information, Fig. S4). This convergence implies that the dynamics are approximately Markovian, and attests to the accuracy of the state-space discretization.

Neither A*β*40 nor A*β*42 exhibits any relaxation process in the *μ*s to ms regime (Fig. 3), in agreement with the findings of the nFCS experiments. ^42^ The slowest relaxation time-scale for A*β*40 is around 85 ns, while for A*β*42 it is around 113 ns. The *t_i_* spectra for both peptides converges on time-scales of ≈ 3 ns. These are the fastest relaxation processes that can be captured by our state-space discretization, and correspond to the local deformation modes of the peptides.

**Figure 3.**
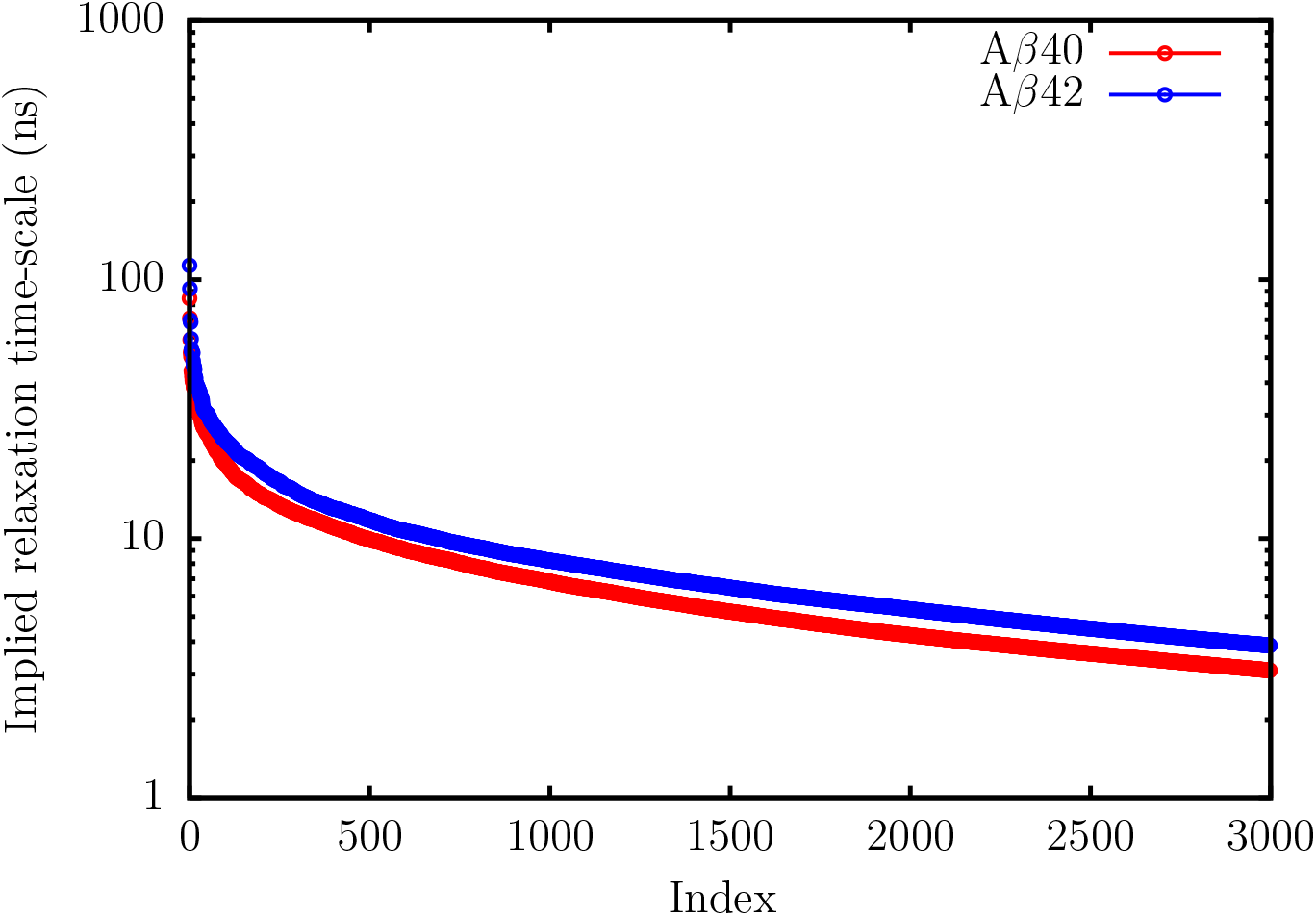
The top three thousand relaxation time-scales, *t_i_*, for A*β*40 (red), and A*β*42 (blue) estimated using Eq. (2). As is evident, there are no relaxation processes in the *μ**s*** to the ms regime. The *t_i_*s start to plateau at around 3ns, and these time-scales are associated with the fastest processes (local deformation modes of the peptides) on the energy landscapes.

### Implication of landscape roughness

The population of fibril-like conformations (N* states) within each free energy basin was computed by using a stringent geometric criterion.^29^ The structural order parameter, 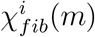, determines the similarity between a conformation *i* within free energy minimum, *m*, and a monomer unit in the experimental fibril structure.

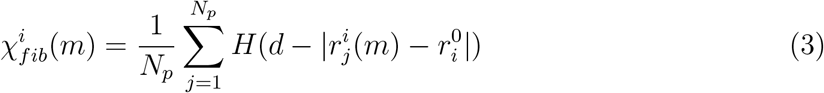

where *H* is the Heaviside step-function; *N_p_* is the total number of pairwise distances; and 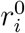 denotes the distance between the *i^th^* pair of beads in the reference structure. For A*β*40, we use the striated fibril structure^40^ as the reference, while for A*β*42, we use the experimental fibril structures corresponding to the U-bend^59^ and the S-bend^41^ morphologies as the reference states. The conformation i is aggregation-prone (assembly-competent) if 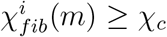. We set *χ_c_* = 0.30 for both A*β*40 and A*β*42. The population of fibril-like conformations within each free energy basin (Fig. 4) is then simply the percentage of constituent structures for which *χ_fib_* ≥ *χ_c_*.

**Figure 4.**
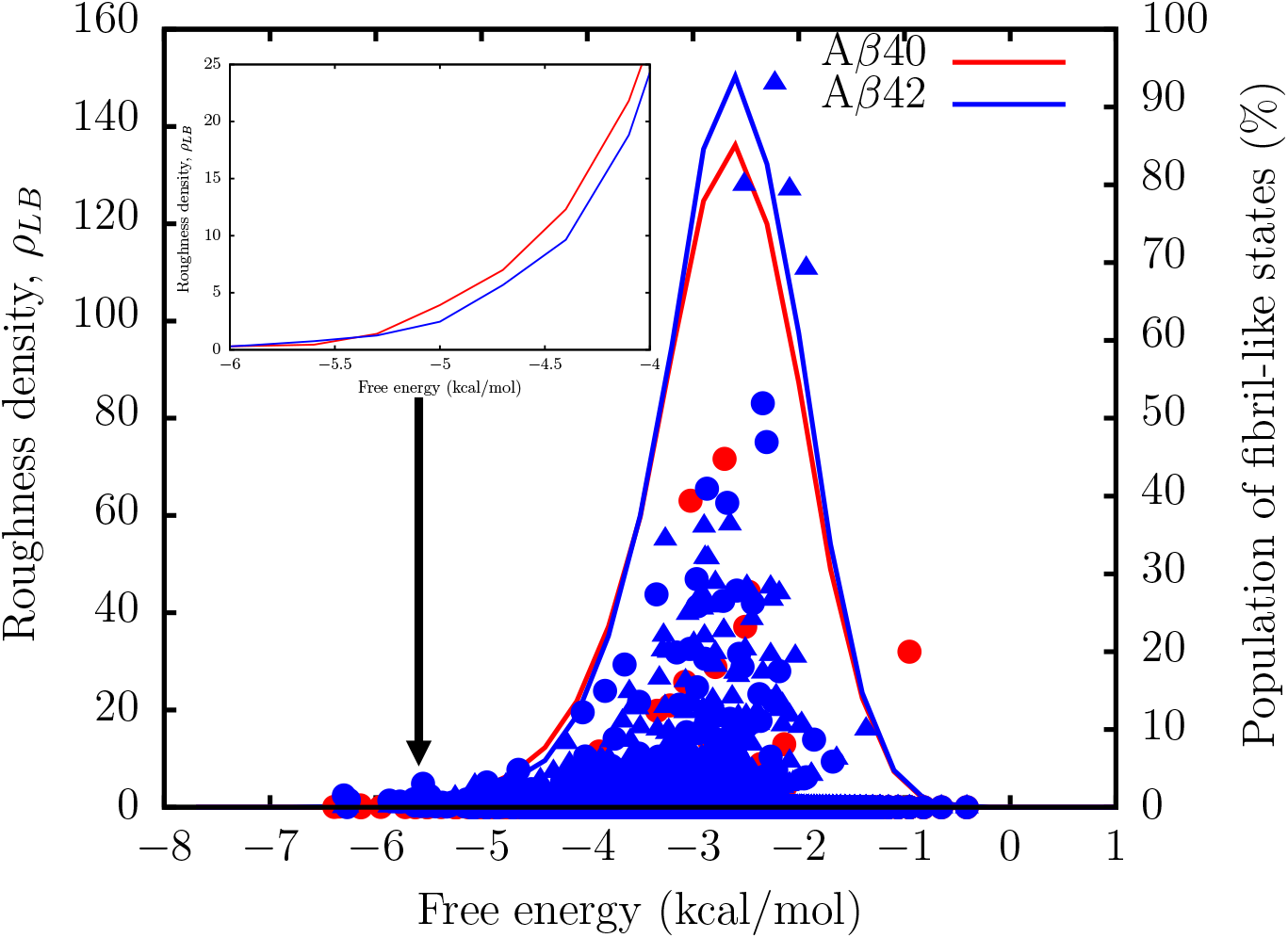
The solid-lines show the variation of the landscape roughness density, *ρ_LB_*, with free energy. The roughness density was computed from the TRDGs using the scheme of Levy and Becker.^66^ The symbols denote the percentages of fibril-like structures within the free energy basins at each energy level: A*β*40 (red circles); A*β*42 U-bend (blue circles); A*β*42 S-bend (blue triangles). Inset: The variation of the landscape roughness for A*β*40 (red) and A*β*42 (blue) close to the free energy of the ground state.

To quantify the ruggedness of the free energy landscape, we computed the roughness density, *ρ_LB_*, from the TRDGs using the formalism of Levy and Becker^66^ (see Supporting Information for details). We find that for both A*β*40 and A*β*42, ρ_LB_, is substantial only over a narrow range of free energies, implying that like other IDPs,^67,68^ the energy landscapes of A*β*40 and A*β*42 exhibit a rather shallow funnel.^66^ This scenario is in contrast to fast-folding globular proteins, where the slope of the energy landscape tends to be quite steep. ^68^

As is evident from Fig. 4, *ρ_LB_* is somewhat higher for the A*β*42 peptide in regions of the landscape where the propensity to find N* states (especially the S-bend motif) is maximal. An immediate implication of this result is that for A*β*42, the landscape in the vicinity of N* states is more undulated compared to A*β*40. Therefore, the aggregation-prone structures have a longer time to form an encounter complex via self-association. Our findings recapitulate the key features of the intramolecular diffusion model for A*β* aggregation kinetics, proposed in a previous work. ^69^

Interestingly, A*β*42 exhibits a smaller *ρ_LB_* close to the free energy ground state (disordered basin) relative to A*β*40 (Fig. 3: Inset). This implies that excitations out of the disordered basin to fibril-like structures is likely to be more favorable in A*β*42.

The different extent of landscape ‘ruggedness’, especially in regions where the probability of finding fibril-like configurations is maximized, appears to be an important feature that distinguishes the statistical ensembles of A*β*42 from A*β*40. Our results suggest that such subtle features at the monomer level could determine the sequence-specific self-assembly propensities.

### N* states are optimal templates for dimerization

To ascertain the self-assembling propensities of different monomer conformations, we probed the dynamics of dimerization, using the number of inter-chain contacts, 〈*N_contacts_*〉, as the order parameter. We assume that a dimer is formed when 〈*N_contacts_*〉 ≥ 4 (see Supporting Information for details). As is evident from the relatively large values of 〈*N_contacts_*〉, dimerization is most effective for S-bend motifs of A*β*42 (Fig. 5). In contrast, the U-bend structures of A*β*42 appear less efficient at templating, with 〈*N_contacts_*〉 attaining large values only at longer time-scales. Dimer formation is the least favored for A*β*40, with fewer inter-chain contacts forming along the trajectories. Hence, the extent of landscape roughness does seem to correlate with the propensity of dimerization.

**Figure 5.**
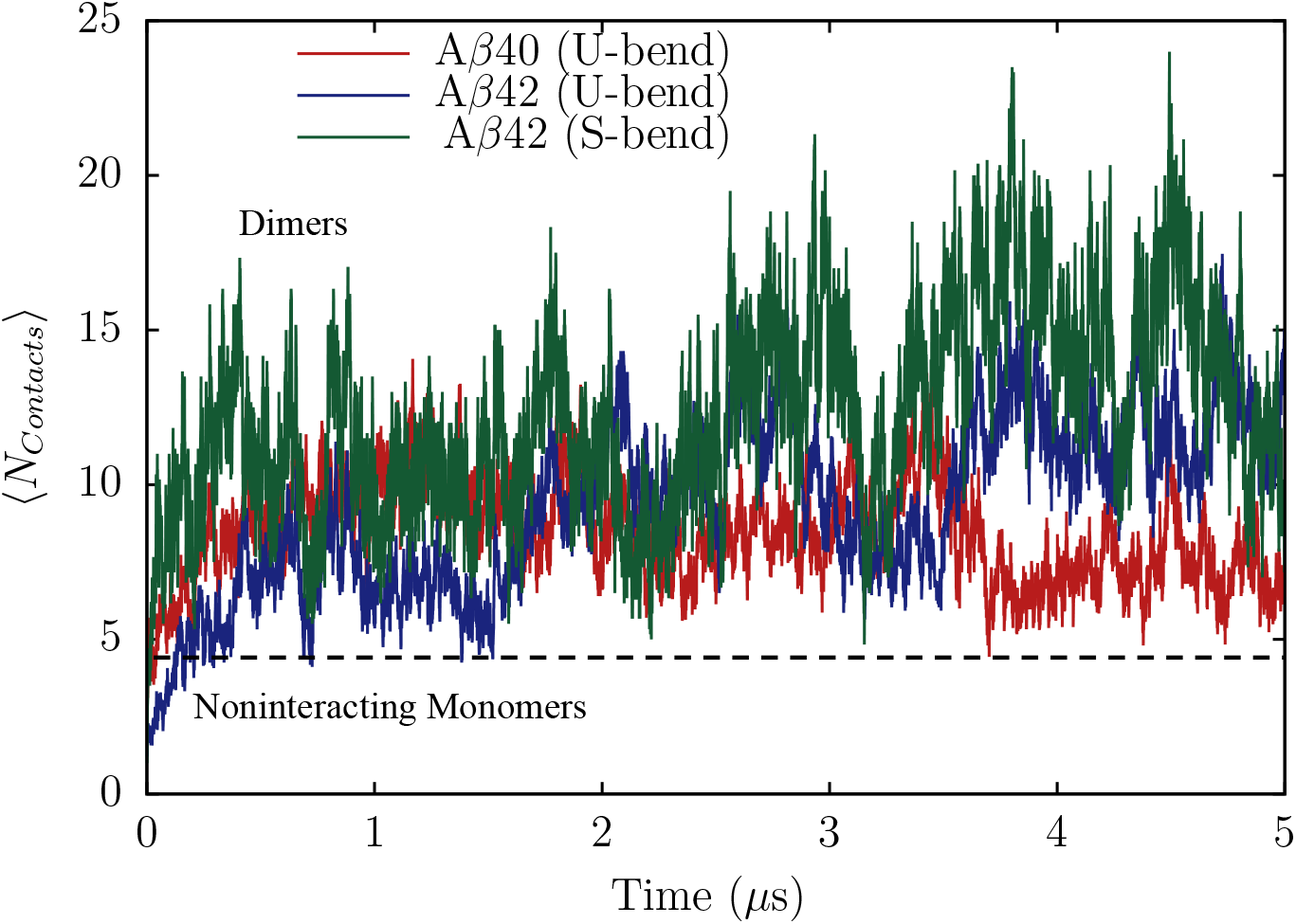
Dimerization kinetics for A*β*40 and A*β*42 when different N* states are allowed to coalesce. The extent of dimerization is monitored using 〈*N_contacts_*〉, the number of inter-chain contacts. Each dimerization profile is generated by averaging the time-dependent changes of *N_contacts_*, over twenty independent trajectories. The dashed line denotes the threshold value of 〈*N_contacts_*〉 for dimer formation.

Interestingly, we observe that dimerization could also occur if one of the monomers is in a N* state, and the other is in a RC-like configuration. Such ‘mixed’ dimers formed by the U-bend structures of A*β*40 and A*β*42 are highly labile, and periodically melt and reform along the trajectories (Supporting Information, Fig. S5). In contrast, mixed dimers formed by the S-bend and RC structures are relatively stable (Supporting Information, Fig. S5). Dimers exhibiting a large number of interchain contacts are also transiently formed through the assembly of U-bend and S-bend conformations of A*β*42 (Supporting Information, Fig. S5). Overall, it appears that the S-bend motif could act as an assembly template for not only N* states, but also for RC-like configurations.

When simulations were initiated from the RC configurations, we observed very little or no dimer formation, suggesting that the free energy barrier for dimerization is much higher for these states (see Supporting Information).

To glean further insight into the dimerization pathways, we constructed transition networks connecting the different substates using hidden Markov models (HMM).^70^ The network for A*β*40 consists of eight states (shown as nodes in Fig. 6). The N* states serve as efficient templates, and dimers in which both the chains have fibril-like signatures form readily. However, these structures relax to thermodynamically more stable configurations within the dimer basin, in which only only one of the chains is fibril-like (Ubend-RC), or both the chains are disordered (random-coil (RC)-dimer). This suggests that the nucleus that will sustain growth exceeds two, and is most likely close to six. ^71^ The connectivity of the transition network suggests that the RC-dimer (lowest energy structure within the dimer basin) is less likely to form through coalescence of two monomeric RC states. In other words, kinetically favored routes for dimerization involve a cascade of free energy excited states, and a direct transition between the monomer and dimer basins seem less efficient.

**Figure 6.**
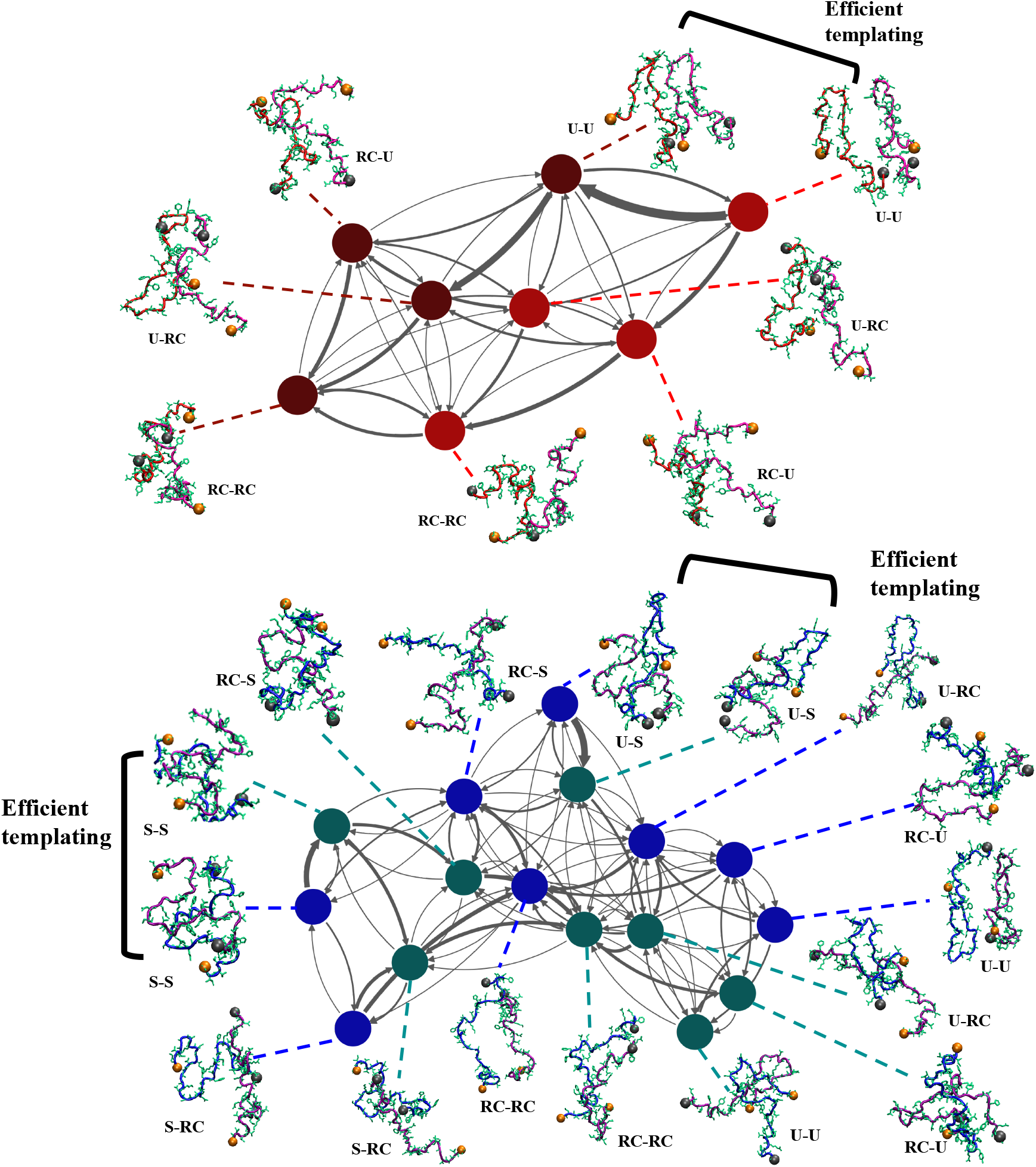
Transition networks illustrating the different dimerization routes for A*β*40 (top) and A*β*42 (bottom). The network was constructed by discretizing the trajectories, and identifying the different substates using a hidden Markov model (HMM). The substates represent the nodes (denoted as circles) of the network. The thickness of the arrows connecting the different nodes are proportional to the transition probabilities. For A*β*40, the nodes denoting the monomers are shown in red, and those denoting the dimers are shown in maroon. For A*β*42, the monomers are represented by blue nodes, and the dimers are represented by teal nodes. Representative snapshots corresponding to each substate are also shown. The N-termini are depicted as orange spheres, and the C-termini are shown as grey spheres. The chains within each substate could either be in a N* (U-bend for A*β*40; U-bend or S-bend for A*β*42) or a random-coil (RC) configuration. Efficient templating implies that the N* state serves as a template for inducing the RC → N* so that dimer formation and other downstream assembly occur readily.

The network for A*β*42 comprises sixteen states, and has a more complex topology (Fig. 6). As is evident from the large transition probability (Fig. 6), the S-bend dimer can form through efficient templating of monomer structures. The mixed dimer unit where one of the chains is in a U-bend configuration, and the other is a S-bend, also assembles rapidly from isolated monomers. These dimers subsequently relax to the more stable RC-dimer through different routes involving multiple intermediates, including the S-bend-RC and the U-bend-RC states.

The connectivity of the network suggest that the U-bend dimer has a low probability of forming via direct coalescence, and once formed it rapidly relaxes to other structures, including the Ubend-RC and the RC-dimer. This equilibration within the dimer basin reflects the growth in 〈*N_contacts_*〉 at long time-scales, when the dimerization reaction is initiated from the U-bend structures of A*β*42 (Fig. 5).

Similar to A*β*40, the probability of a direct transition between the ground states of the monomer and the dimer is low. For dimerization to be efficient, the pathway must proceed through free energy excited states within the different basins, thus supporting the expectations based on the N* theory.

### Conformational transitions to fibril-like states and implications of Ostwald’s rule

We calculated the kinetics associated with the transition from the free energy ground state (RC-like structures) to N* states (with *χ_fib_* ≥ *χ_c_*) using the distributions of first passage times (FPTs), *P*(*τ_FPT_*), where:

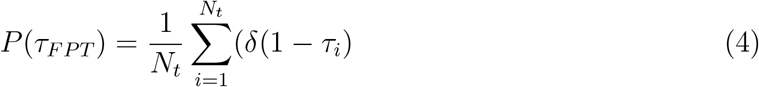

In Eq. (4), *τ_i_* is the time in the *i^th^* trajectory when a structure with *χ_fib_* ≥ *χ_c_* is visited for the first time, and *N_t_* denotes the total number of independent trajectories.

The distributions of the FPTs are shown in Fig. 7 and Fig. 8. The widths of *P*(*τ_FPT_*)s imply that the transitions could occur through a multitude of pathways. Interestingly, all the distributions are approximately Poissonian, suggesting that the transitions to the N* states do not involve any long-lived intermediates, and the underlying kinetics can be appropriately described by a two-state model.

**Figure 7.**
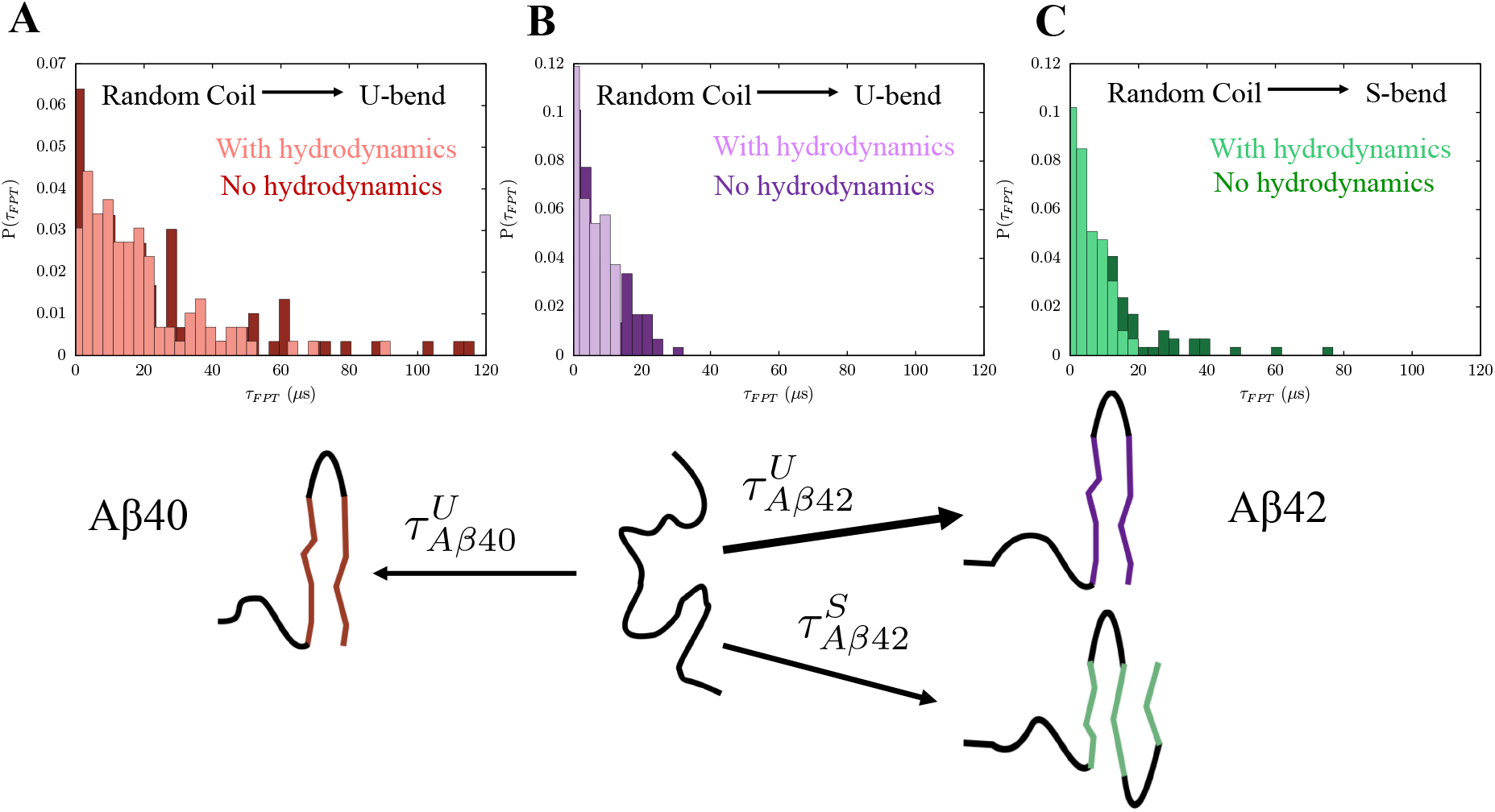
Histograms depicting the distributions of first passage times (FPTs) corresponding to the transition from the disordered ground state to the fibril-like N* states. A schematic describing the different transitions is shown below the histograms. (A) The FPT distribution for the transition from an equilibrium RC (free energy ground state) to the U-bend fibril-like structures in A*β*40. In the absence of hydrodynamic interactions, the MFPT, 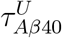, for this transition is ≈ 25 *μ*s. (B) The FPT distribution for the transition from an equilibrium RC to the U-bend fibril-like conformations in A*β*42. In the absence of hydrodynamic interactions, the MFPT,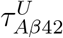, for this transition is ≈ 9 *μ*s. (C) The FPT distribution for the transition from RC-like structures to the S-bend fibril-like conformations. The MFPT, 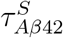 larger than 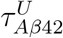, and is around 13 *μ*s. In all cases, inclusion of hydrodynamic interactions shortens the tails of the FPT distributions, and accelerates the conformational transitions to the N* states.

**Figure 8.**
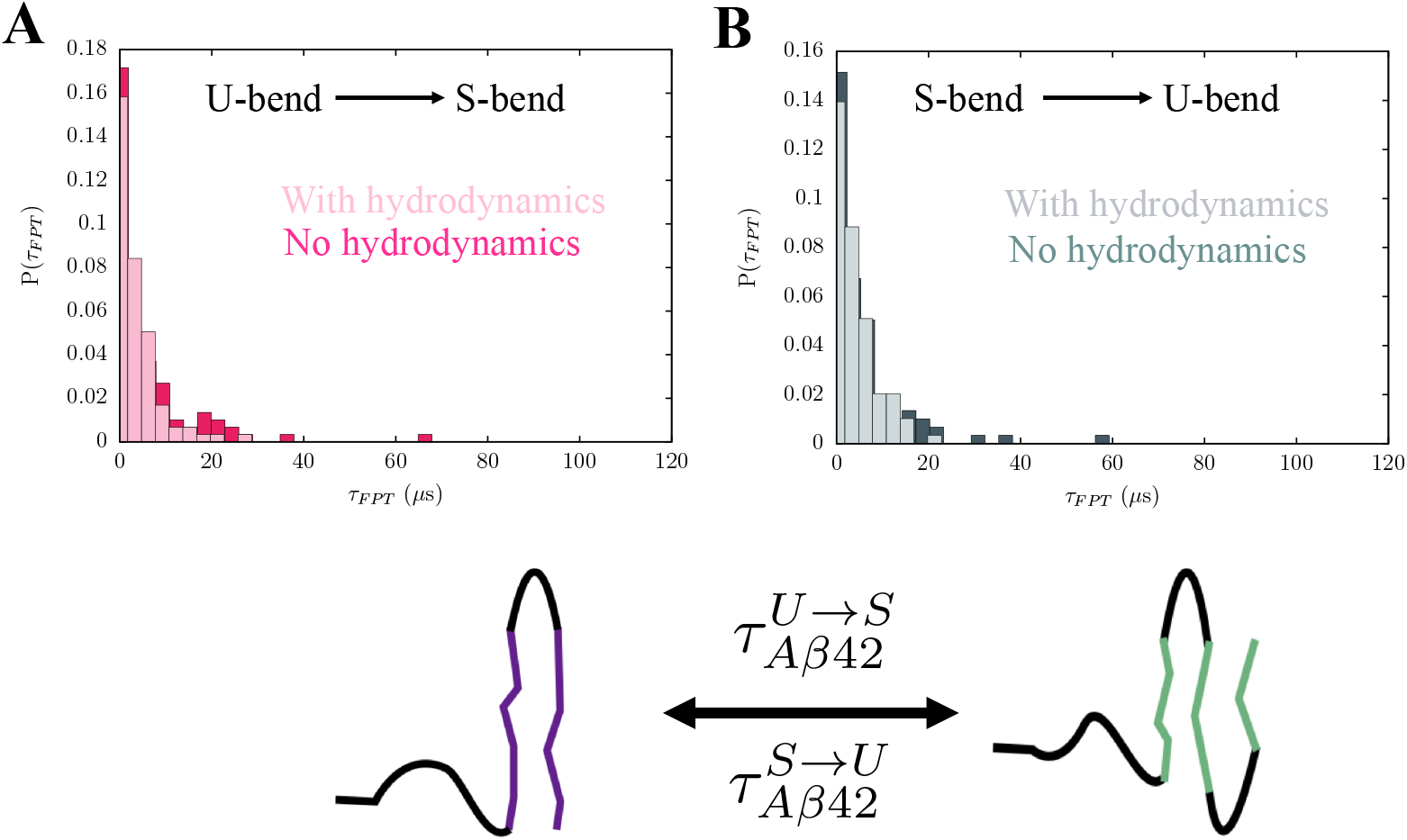
The FPT distributions corresponding to the conformational switch between the U-bend and S-bend fibril-like conformations in the ensemble of A*β*42. A: FPT distribution for the U-bend to S-bend transition. B: FPT distribution for the S-bend to U-bend transition. In both the cases, the MFPT is ≈ 7 *μ*s. The FPT distributions do not exhibit long tails when hydrodynamic interactions are taken into account. The MFPTs corresponding to the conformational switch also become smaller.

The transitions to fibril-like monomer configurations are considerably faster in A*β*42 as compared to A*β*40 (Fig. 7). The mean first passage time (MFPT), 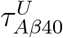, associated with the transition from an equilibrium RC-like conformation to a U-bend structure in A*β*40 is ≈ 25 *μ*s. The corresponding MFPT for A*β*42, 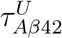 is around three times smaller (≈ 9 *μ*s). Although the S-bend structure is thermodynamically preferred,^29^ it forms on a longer time-scale as compared to the U-bend conformation 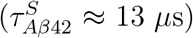.

The ordering kinetics of polymorphic structures in A*β*42 is in accord with Ostwald’s rule of stages. ^72^ The thermodynamically favored S-bend forms at a slower rate than the less stable U-bend topology. Ostwald’s rule was originally proposed in the context of crystal polymorphism in materials (graphite and diamond structures in carbon). However, it’s seem to be far-reaching, and as illustrated here, could encompass disorder-to-order transitions in IDPs.

The conformational switch between the U-bend and S-bend forms in A*β*42 is faster than the corresponding RC → N* transitions, with 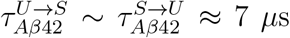 (Fig.8). This observation implies that for a majority of the conformational switching events, neither the U- or the S-bend structures have to access the RC state. Overall, the MFPTs for the different conformational transitions in A*β* peptides are in good agreement with the recent predictions from all-atom simulations. ^39,73^

#### Effect of Hydrodynamic Interactions

We find that the inclusion of hydrodynamic interactions not only accelerates the disorder to order (RC → N*) transitions, but also the conformational switch between the U-bend and S-bend motifs of A*β*42. Interestingly, the long-tails of the FPT distributions, which arise due to an ensemble of non-optimal transition paths, become less pronounced in the presence of hydrodynamic interactions. Our observations are in accord with recent works that underscore the importance of hydrodynamic interactions in guiding the collapse transitions in proteins, ^74^ synthetic polymers, ^75^ and stepping of molecular motors on microtubule tracks. ^76^

For most transitions, there is a ≈ 1.3-1.5 fold decrease in MFPT in the presence of hydrodynamic interactions (Supporting Information, Table S3). However, hydrodynamic interactions have a more dominant impact on the RC → S-bend transition (Fig. 7(C)), with the MFPT being reduced by a factor of ≈ 2. We speculate that this enhancement of the reaction rate in the case of the S-bend motif could be linked to the more complex molecular mechanism (as compared to U-bend structures) underlying its formation.

## Discussion

In this work, we used the SOP-IDP model^29^ to characterize the structural, as well as kinetic heterogeneity of A*β*40 and A*β*42 monomers. Structural heterogeneity, which is evident at the monomer level, has recently been illustrated to play an important role in the aggregation cascade in insightful single-molecule fluorescence imaging experiments.^11^ The topography of the free energy landscapes for both the sequences suggest that the RC-like ground state is readily accessible from the N*-like states found in the fibril polymorphs, over a wide range of experimental conditions, such as temperature, pH or salt concentration. Hence, it is not surprising that the RC-like conformations belonging to the ground state determine the different thermodynamic observables, including residue-residue contact maps, secondary structure profiles, and residue-dependent chemical shifts.

The dominance of the featureless ensemble of the ground state might give the erroneous impression that not much can be discerned from the study of monomers. However, we find that monomer conformations, which have remnants of the fibril state, are encoded as excitations in the free energy landscape (Fig. 9). In other words, the “ordered” structures appear as high-lying free energy minima. Such a topography seems consistent with the recently proposed ‘inverted funnel’ free energy landscape picture proposed for IDPs.^77^

**Figure 9.**
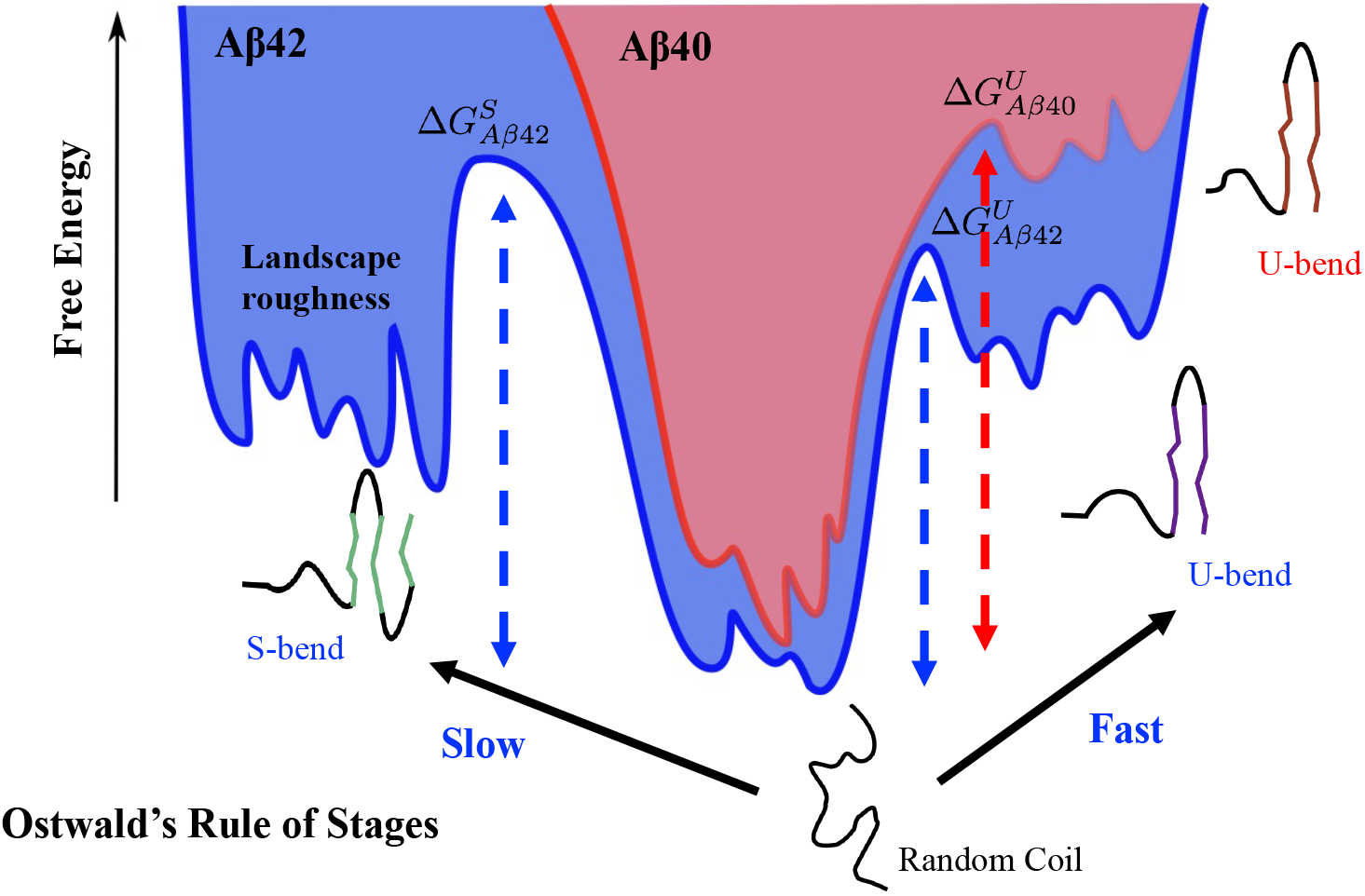
A schematic of the free energy landscape for the A*β*40 (red) and A*β*42 (blue). The landscape roughness is enhanced in A*β*42 (especially near the S-bend motif) compared to A*β*40. This subtle variation in topography could have important implications on the self-assembling propensities of the monomers. Also shown are the relative positions of the different N* states relative to the disordered ensemble (free energy ground state for both A*β*40 and A*β*42). As is evident from the relatively high free energy barrier,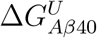, the transition to the U-bend form is slower in case of A*β*40. The transitions to the fibril-like conformations occur much faster in A*β*42, and is dictated by Ostwald’s rule of stages. The U-bend topology appears before the S-bend topology (i.e. 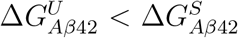) because the later is thermodynamically favored.

### Relaxation kinetics in the free energy landscape

Time-resolved spectroscopy has emerged as a potent tool for the characterization of the multi-tiered conformational dynamics in IDPs. ^42,43,69^ Nanosecond fluorescence spectroscopy (nFCS) showed that the reconfiguration time of A*β*40 and A*β*42 monomers is ≈ 35 ns,^42^ and neither peptide exhibits conformational dynamics on the *μ*s to ms time-scale. On the other hand, based on Trp-Cys contact quenching experiments, were used to estimate an upper-bound of around 1 *μ*s for the intramolecular diffusion of A*β* peptides. ^69^ In contrast to global chain relaxation, localized motions of the peptide backbone, including chain-tumbling, and segmental dynamics occur on much faster time-scales (≈ 10 ns or less) and have recently been reported using NMR spin-relaxation techniques. ^43^

The dynamics of A*β* monomers are hierarchically organized, with the relaxation times ranging from ≈ 3 ns to 100 ns. The slower time-scales correspond to global chain relaxation, while the faster ones correspond to local deformation modes of the peptide chains. In accord with nFCS experiments, ^42^ we do not find any relaxation process in the *μ*s to ms regime. Some studies^78–80^ speculate that certain IDP sequences may exhibit glass-like behavior, and switch forms over extremely long time-scales. However, no such behavior for the A*β* peptides, is immediately apparent from the organization of the energy landscapes, and relaxation time-scales (Figs 1 and 2). It is likely that the length of the IDP has to be sufficiently enough to see the manifestation of glassy dynamics. ^80^

### Landscape ruggedness peaks near N* states

Despite having globally similar topographies, the landscape ruggedness of A*β*40 and A*β*42 is different in the regions with high N* populations. We believe that this does have important implications in dictating the distinct aggregation propensities of A*β*40 and A*β*42. The enhanced ruggedness impedes intramolecular diffusion in case of A*β*42, thus prolonging the lifetime of the N* states, which greatly facilitates self-assembly events. Because ruggedness decreases the diffusion by *e*^(*δϵ/k_B_T*)2^ (where δ*e/k_B_T* is the roughness scale, even a small change in δe could have a large effect on diffusion, ^81^ and hence the lifetime of the N* state.

### Ostwald’s rule and structural polymorphism

The conformational transition from the disordered state to fibril-like structures in the monomers occurs on the *μ*s time-scale. The transition to the fibril-like monomer state is faster in A*β*42 compared to A*β*40. This observation implies that A*β*42 is better poised to form the early oligomers along the aggregation cascade. ^82,83^

Surprisingly, we find that in A*β*42, the transition rates to the U-bend and S-bend structures are in accord with Ostwald’s rule of stages, which is often used in the solid-state community to predict the appearance of different crystal polymorphs. In other words, the U-bend fibril polymorph is likely to form first, while the S-bend polymorph, which is thermodynamically more stable, should appear on longer observation time-scales. Of course, the precise order of events will be dictated by the underlying energy landscape, which can be tilted by changes in external conditions.

As shown here, Ostwald’s rule is manifested even during the early stages of the disorder to order transition, in the monomer conformational ensemble. However, it could also have important consequences during the late events of aggregation. A recent study has shown that Ostwald’s rule dictates the structural transitions in dipeptide supramolecular polymers. ^72^ These findings could be important to advance our understanding of fibril polymorphism, particularly for sequences with low-complexity (LC) domains,^84–86^ where even subtle variations in experimental conditions, or preparation protocols result in different fibril morphologies.

It is worth pointing out that Ostwald’s rule also explains tidily the ordering kinetics in the low-complexity domain fused in sarcoma (FUS) protein and various related constructs. In the FUS-LC sequence (residues 1-214), only residues 39-95 form S-bend type fibrils (core1). On the other hand, residues 112-150 in a truncated variant of FUS-LC (residues 108-214), forms a fibril with a U-bend topology (core2), which is marginally destabilized with respect to core1. In an intriguing new development, ^87^ it has been shown that another C-terminal variant of FUS-LC (residues 141-214) could also forms fibrils (core3), with residues 155-190 forming the core region. Experiments suggest that core3 forms before core2, which in turn forms before the core1. The stabilities of the three cores show exactly the opposite trend.^87^ Thus, there is an inverse relationship between thermodynamic stability and fibril formation rates, further affirming Ostwald’s rule of stages, that we have established in this study in the context of A*β* aggregation.

### N* theory for Huntington (htt) polyglutamine (Q)

The theory based on the N* provides support to the kinetic scheme developed for dimer formation from the disordered ground state of the amphiphilic domain, Q_7_.^88^ Using relaxation dispersion NMR measurements it was shown that the ground state of Q_7_ is disordered (population ≈ 95%), just as in A*β* peptides. The productive route to Q7 dimerization occurs with small probability (≈ 2%) on a time scale of about 20μs. The unstable dimers coalesce to form stable tetramers. The detailed quantitative kinetic analysis presented in the NMR study^88^ is in accord with the N* theory elucidated here. In accord with the mechanism for A*β* dimer formation, we envision that the most productive path must involve 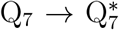, which is followed by 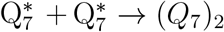. From this picture, it follows that 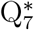 would be the sparsely populated (< 2%) excited state in the monomer ensemble. The accumulating experimental evidence (see for example^30,88,89^) backed up by computations show that the N* theory is a general mechanism in the initiation of protein aggregation. A corollary is that characterization of excited states is necessary to assess the aggregation propensity of protein sequences.

## Concluding Remarks

A*β*40 and A*β*42 are the major isoforms implicated in Alzheimer’s disease, a progressive neurodegenerative disorder that affects a large fraction of the global population. Despite being present in relatively lower concentrations, A*β*42 aggregates nearly an order of magnitude faster than A*β*40.^82^ For both the peptides, many experimentally relevant thermodynamic observables, such as *R_g_* and FRET efficiencies, are solely determined by the free energy ground state (consisting of random-coil (RC) like structures). Hence, at the monomer level, a description based on ensemble-averages alone is inadequate for rationalizing the apparent anomaly in the aggregation behavior of A*β*40 and A*β*42. In this study, we show that a detailed view of the free energy landscape (and not simply the thermodynamic ground state) is necessary for deciphering alloform-specific differences. The aggregation-prone, fibril-like monomer conformations (N* states) appear as excitations on the energy landscape, and can be transiently accessed from the RC configurations on the *μ*s time-scale. The *RC* → *N*\* transition is several times faster for A*β*42, implying that it is kinetically predisposed to assemble compared to A*β*40. Surprisingly, we find that for A*β*42, the least stable fibril-like structure (U-bend) forms faster than more stable ones (S-bend), in accord with Ostwald’s rule of stages, ^72^ which was postulated nearly a century ago in the context of crystal polymorphism.

We show that the extent of landscape roughness, particularly in regions where assembly-like conformations are most likely to be found, tidily explains the oligomerization propensity of A*β*40 and A*β*42. In particular, we find that the S-bend configurations, which are exclusively found in the conformational ensemble of A*β*42, act as very efficient templates of self-assembly. To probe the multitude of dimerization pathways, we constructed transition networks from our simulations using the formalism of hidden Markov Models (HMMs). ^70^ We find that kinetically favorable dimerization routes proceed through N* states, although the complex topology of the networks suggest that other possibilities could also exist. It is likely that dimerization, and by inference, higher-order oligomerization pathways, are modulated by external conditions (such as pH, presence of crowders or denaturants),^90^ or by interactions with membranes. ^91^

## Computational Methodology

### The SOP-IDP Model

The simulations of the A*β*40 and A*β*42 peptides were carried out using the recently introduced Self-Organized Polymer (SOP) model for intrinsically disordered proteins (abbreviated as SOP-IDP). ^29,92^ In this model, each amino acid residue is represented using two interaction sites: a backbone bead (BB) centered on the *C_α_* atom, and a side-chain (SC) bead centered on the center-of-mass of the side chain (Fig. S1). The SOP-IDP energy function is given by:

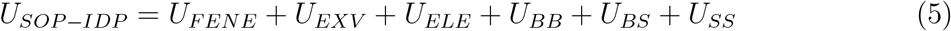

In Eq. 5, *U_FENE_* describes the chain connectivity between the different interaction sites; *U_EXV_* denotes purely repulsive interactions, which prevent any unphysical overlap between beads; *U_ELE_* describes the electrostatic interactions between charged amino-acid side chains. The final three terms, *U_BB_, U_BS_* and *U_SS_* are Lennard-Jones type potentials, which describe the backbone-backbone (BB), backbone-side-chain (BS) and side chain-side chain (SS), respectively. Importantly, *U_SS_* encodes the sequence-specificity of the model, and is based on the Betancourt-Thirumalai interaction map.^93^ The detailed functional form of the SOP-IDP potential, and the force-field parameters are included in the Supporting Information.

### Monomer Simulations

To probe the conformational dynamics of A*β* monomers, we carried out Brownian dynamics (BD) simulations in the high friction regime, corresponding to a solvent viscosity of 10^-3^ Pa.s. The inertial term in the Langevin equation can be ignored in this limit, and the motion of each bead *i* is described by:

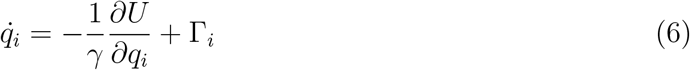

In the over-damped limit, the natural unit of time 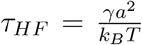. The typical values of the energy and length scales are 1kcal/mol and 1Å, respectively. Using these values, *τ_HF_*, is estimated to be 13.2 ps. The equations of motion were integrated using the Ermak-McCammon algorithm,^94^ employing a time-step of 0.05*τ_HF_*. To obtain meaningful estimates of first passage times (FPTs), and other kinetic observables, we carried 100 independent simulations for each A*β* monomer at 298 K. Each trajectory consisted of 2 × 10^7^ steps.

### Dimerization simulations

We probed the kinetics of dimerization in A*β*40 and A*β*42 using BD simulations. To mimic the critical protein concentration required for dimerization, we followed the setup described previously,^95^ byconstraining the distance between the centers of masses of the two A*β* chains to the thermally averaged *R_g_* of the monomer. The nonbonded interactions among beads in different chains were modeled as if they were part of the same monomer. In other words, if two residues *R*_1_ and *R*_2_ within the monomer have a collision diameter, *σ*_12_, and an interaction strength, *ϵ*_12_, then *R*_1_ and *R*_2_ in different A*β* chains would also interact with the same energy function.

The dimerization process was initiated from configurations where both A*β* chains adopted a N* (fibril-like) conformation (U-bend for A*β*40; U-bend or S-bend for A*β*42), only one of the chains adopted a N* conformation, both chains were in a random-coil (RC) configuration. For each initial condition, we carried out 30 independent simulations of 8 × 10^6^ steps, corresponding to ≈ 5.3,*μ*s. The extent of dimerization was quantified using *N_contacts_*, the number of inter-chain contacts formed along the trajectory. An inter-chain contact was assumed to form if the distance between any two beads on different A*β* chains was ≤ 6 Å.

### Modeling Hydrodynamic Interactions

To simulate the effect of hydrodynamic interactions, we carried out BD simulations, where the motion of each bead is described by:

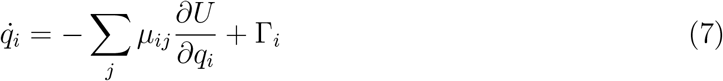

In Eq. 3, *μ_ij_* denotes the conformation-dependent mobility tensor, and is computed using a modified form of the Rotne-Pragar-Yamakawa approximation^96^ introduced by Zuk et al.^97^

### Constructing Monomer Free Energy Landscapes

The conformational ensembles obtained from the simulations were clustered into kinetically distinct microstates using the distribution of reciprocal interatomic distance (DRID) metric.^44^ Each microstate *i* can be considered to be a local minimum on the free energy landscape. The free energy of minimum i was estimated as *F_i_* = –*k_B_T* ln(*N_i_*), where *N_i_* denotes the number of conformations in cluster *i*. ^54^ The apparent free energy barrier between the different minima were estimated using the ‘minimum-cut’ procedure proposed by Krivov and Karplus.^47,98^ The number of transitions, nj between minima i and j were converted to symmetrized edge-capacities, *c_ij_* = (*n_ij_* + *n_ji_*)/2. Within this framework, the free energy landscape can be considered as a flow-network having capacitated edges,,^98^ with the nodes denoting the different local minima. The free energy barrier between minima can be estimated by computing the number of minimum cuts, *N_ij_*., between the nodes in a pairwise fashion. Here, we used the Gomory-Hu procedure to transform the flow-network into a tree, and compute *N_ij_*. The effective barrier between minimum *i* and *j* is given by *F_ij_* = –*k_B_T* ln(*N_ij_*).

The free energy landscapes were visualized in the form of transition disconnectivity graphs (TRDGs).^45,48^ The TRDG representation provides a faithful description of the underlying kinetics, in contrast to many schemes based on low-dimensional projections of the energy landscape, ^47^ where basins separated by high free energy barriers may be lumped together. ^99^ The transition disconnectivity graphs (TRDGs) were constructed using the disconnectionDPS code.^100^ Further details can be found in the Supporting Information.

### Constructing Transition Networks for Dimerization

Each snapshot in the dimerization trajectories were assigned a binary code (with values being either 0 or 1) based on the number of inter-chain contacts, *N_contacts_*, and the structural overlap, *χ_fib_*(*t*), with the monomer unit of the experimentally determined fibril structures. For A*β*40, the snapshots were indexed as 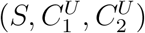, with *S*=1 denoting a dimer (*N_contacts_* ≥ 4) and *S*=0 denoting isolated monomers. The label 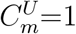 if the chain, *C_m_*, (*m* is either 1 or 2) is fibril-like (*χ_fib_*(*t*) ≥ 0.30) and 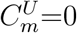 if the chain adopts a RC-like configuration.

For A*β*42, each configuration has two additional labels due to the possibility of the S-bend configuration. The snapshots in the trajectories were indexed as 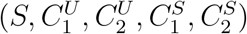. In this case, *S*=1 represents a dimer, and *S*=0 denotes isolated monomers. The label 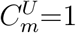 if the chain, *C_m_*, (*m* is either 1 or 2) adopts a U-bend configuration (*χ_fib_*(*t*) ≥ 0.30) and 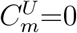 it the chain is RC-like. Similarly, 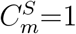 if the chain, C_m_, adopts a S-bend configuration, and 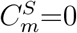 if the chain is in a RC-like configuration.

After assigning a discrete label to each snapshot, transition networks describing the dimerization process for A*β*40 and A*β*42 were constructed using the hidden Markov Model (HMM) formalism^70^ implemented within the *PyEMMA2.5* distribution.^101^ The HMMs were estimated at a series of lag-times, and to construct the transition networks we chose a lag-time where the implied time-scales appear converged (Fig. S6).

## Supporting Information

### The SOP-IDP model

The Self-Organized Polymer (SOP) model for intrinsically disordered models (abbreviated as SOP-IDP) was used to simulate the conformational dynamics of the A*β*40 and A*β*42 monomers. The SOP-IDP model quantitatively reproduces the scattering profiles of a diverse range of IDP sequences of varying sequence composition, lengths, charge densities.^1^ In the SOP-IDP model, each amino acid residue is represented using two interaction sites: a backbone bead (BB) centered on the *C_α_* atom, and a side-chain bead (SC) centered on the center-of-mass of the side-chain (Fig. S1). The energy function for the SOP-IDP model is given by:

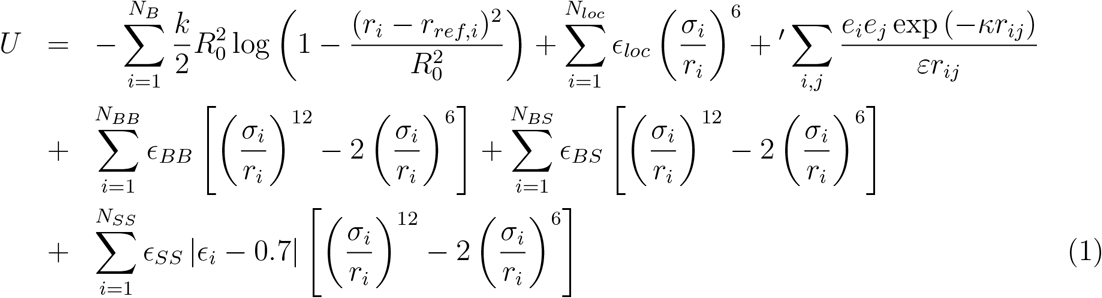

The first term in Eq. 1 denotes the finitely extensible nonlinear elastic (FENE) potential, which accounts for the chain connectivity, with *r_ref,i_* representing the equilibrium distance between the bonded moieties. Purely repulsive excluded volume interactions are included to (second term in Eq. 1) prevent any unphysical overlap between the beads. The third term describes the Debye-Hückel potential, which accounts for the electrostatic interactions between the different charged residues. The final three terms in Eq. 1 describe the backbone-backbone (BB), backbone-sidechain (BS), and sidechain-sidechain (SS) interactions, respectively. The parameter *ϵ_i_* corresponds to the Betancourt-Thirumalai matrix element,^2^ and encodes the sequence-specificity of the SOP-IDP model. There are three adjustable parameters in the model: *ϵ_BB_*, *ϵ_BS_*, and *ϵ_SS_*. Their values were determined using a learning procedure described elsewhere.^1^ The values of the different parameters used in the SOP-IDP model are given in Tables S1 and S2.

**Figure S1.**
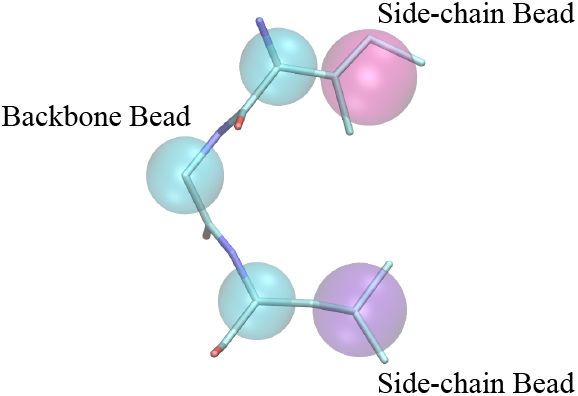
A schematic illustrating the coarse-grained representation used in the SOP-IDP model, where each amino-acid is represented by two beads (interaction sites). One bead is centered on the backbone C_α_ atom, and the other one is centered on the center-of-mass of the side-chain. Here, the side-chain beads are shown in different colors to illustrate the sequence-specificity encoded in the SOP-IDP model.This figure is adapted from ref.^3^

**Table S1:** Energy function parameters for the SOP-IDP model.

| Parameter | Value |
| --- | --- |
| $R_0$ | 2.0 Å |
| $k$ | 20.0 kcal/(mol Å<sup>2</sup>) |
| $\epsilon_{loc}$ | 1.0 kcal/mol |
| $\epsilon_{BB}$ | 0.12 kcal/mol |
| $\epsilon_{BS}$ | 0.24 kcal/mol |
| $\epsilon_{SS}$ | 0.18 kcal/mol |

**Table S2:** Parameters for the coarse-grained beads used in the SOP-IDP model.

| Residue | Radius (Å) | charge (e) |
| --- | --- | --- |
| Gly | 0.0 | 0.0 |
| Ala | 2.52 | 0.0 |
| Val | 2.93 | 0.0 |
| Leu | 3.09 | 0.0 |
| Ile | 3.09 | 0.0 |
| Met | 3.09 | 0.0 |
| Phe | 3.18 | 0.0 |
| Pro | 2.78 | 0.0 |
| Ser | 2.59 | 0.0 |
| Thr | 2.81 | 0.0 |
| Asn | 2.84 | 0.0 |
| Gln | 3.01 | 0.0 |
| Tyr | 3.23 | 0.0 |
| Trp | 3.39 | 0.0 |
| Asp | 2.79 | -1.0 |
| Glu | 2.96 | -1.0 |
| His | 3.04 | 0.0 |
| Lys | 3.18 | 1.0 |
| Arg | 3.28 | 1.0 |
| Cys | 2.74 | 0.0 |
| backbone | 1.90 | 0.0 |

### Identifying free energy local minima

The conformations sampled by the BD trajectories was partitioned into discrete microstates using structural clustering. First, each conformation was mapped into its equivalent distribution of reciprocal interatomic distance (DRID) metric.^4^ As shown by Zhou and Calfisch,^4^

**Table S3:** A comparison of the mean first passage times (MFPTs) for the various transitions obtained with hydrodynamic interactions (*τ_HD_*) and without hydrodynamic interactions (*τ_NHD_*).

| Transition | $\tau_{HD}$ (μs) | $\tau_{NHD}$ (μs) |
| --- | --- | --- |
| RC → U-bend (Aβ40) | 19 | 25 |
| RC → U-bend (Aβ42) | 6 | 9 |
| RC → S-bend (Aβ42) | 7 | 13 |
| U-bend → S-bend (Aβ42) | 5 | 7 |
| S-bend → U-bend (Aβ42) | 5 | 7 |

DRID clustering usually preserves the kinetic distances between any two given conformations, and is hence better suited for kinetic analysis as compared to other conventional structurebased metrics. In order to compute the DRID, two sets of atoms are required: the first is a set of *m* centroids, and the other is a list of atoms, *N_atoms_*. Each centroid, *i*, is associated with the three moments of the distribution of reciprocal distances (*μ_i_, v_i_, ζ_i_*), which describe the structural features of a given conformation. Hence, each conformation is described by a DRID vector of dimension 3*m*. The distance, *s_jk_*, between any two conformations *j* and *k* in the space of DRID vectors is given as:

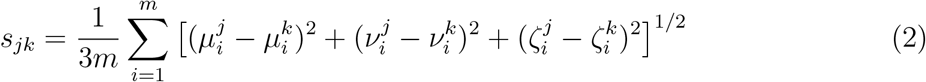

The moments of distribution of the reciprocal distances are expressed as:

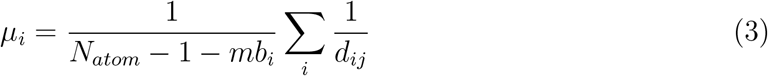

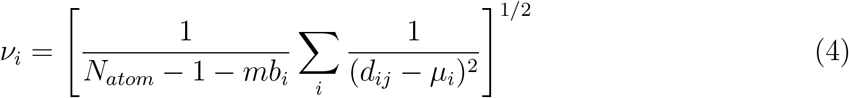

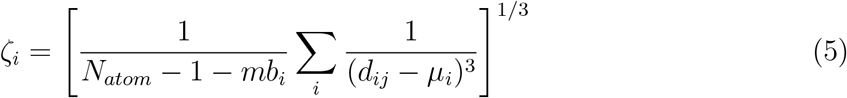

In the equations above, *d_ij_* is the distance of atom *j* from centroid *i*; *N_atom_* is the number of atoms; *nb_i_* is the number of atoms bonded to centroid *i*. The summations in Eq. 1-3 do not include the centroid *i* and the atoms covalently linked to it.

Subsequent to mapping in the DRID space, the conformational ensemble was clustered using a regular space clustering algorithm implemented within the *PyEMMA2.5* distribution.^5^ To identify distinct microstates, a cutoff of 0.15 was used for *s_jk_*. Using this procedure, we identified 5956 microstates for A*β*40, and 6639 microstates for A*β*42. Each microstate *i* can be considered to be a local minimum on the free energy landscape. The free energy of minimum i was estimated as *F_i_* = – *k_B_T* ln(*N_i_*), where *N_i_* denotes the number of conformations in cluster *i*.^6^

**Figure S2.**
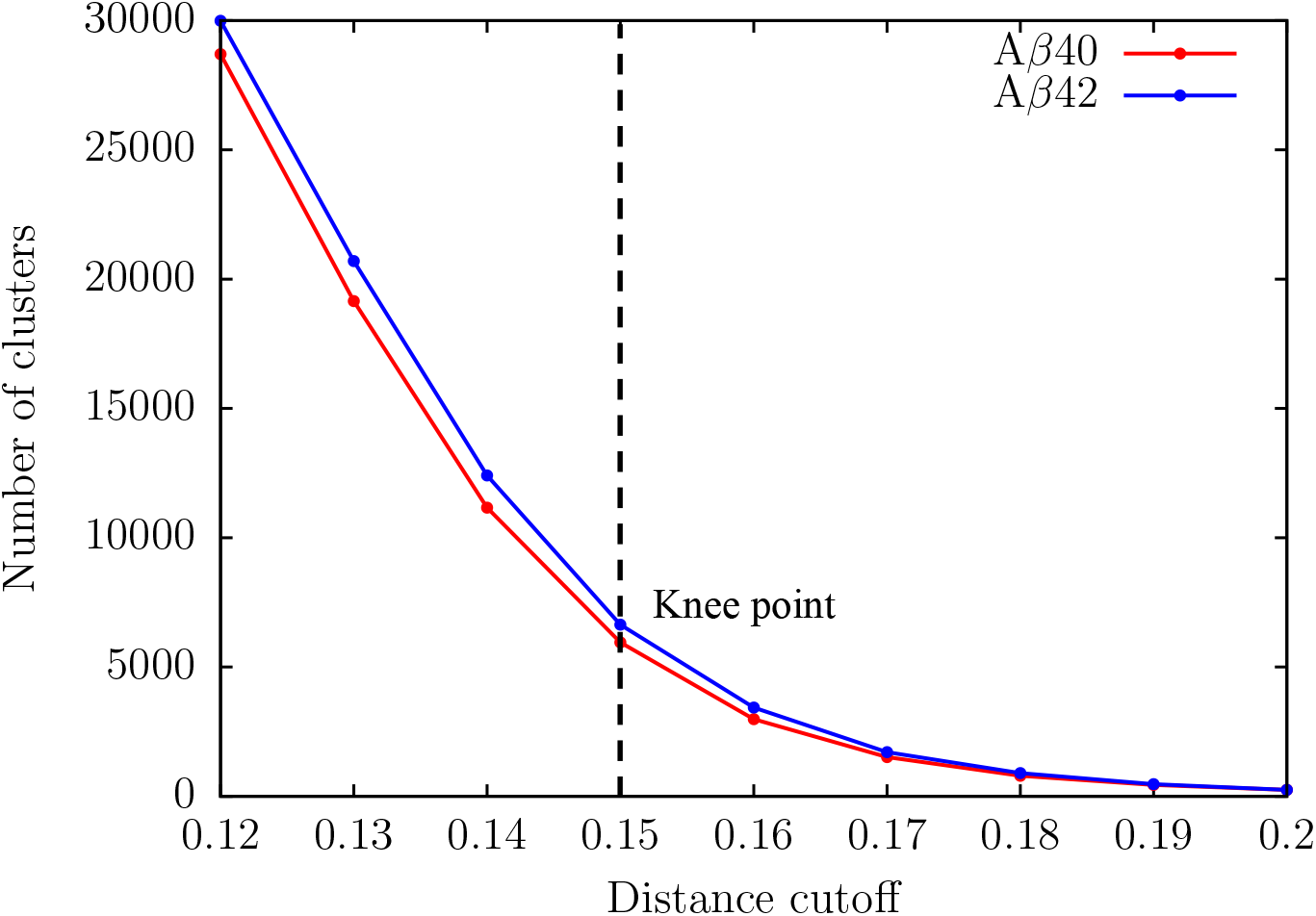
Variation of the number of clusters with distance cutoff for A*β*40 (red) and A*β*42 (blue). The optimal distance cutoff in each case was determined from the knee point (denoted as a dashed line).

### Estimating effective barrier heights

The number of transitions, *n_ij_*, between cluster *i* and *j* (or equivalently, the free energy local minima), can be directly computed from the BD trajectories. These counts were converted to symmetrized edge capacities, *c_ij_* = (*n_ij_* + *n_ji_*)/2 following Krivov and Karplus^7^ The edge-capacities reflect the underlying dynamics of the system. Based on the isomorphism between the free energy landscape, and a flow-network consisting of capacitated edges, calculating the effective barrier between two minima is equivalent to determining the maximum flow in the network.^8^ According to the Ford and Fulkerson theorem,^9^ the maximum flow (i.e. effective free energy barrier) between two nodes (or local minima) can be obtained from the number of “minimum-cuts”. Instead of finding the number of minimum cuts, *k*, for each pair of minima separately, we use the Gomory-Hu procedure^10^ as implemented within the *networkx* module. The Gomory-Hu procedure converts the flow-network with capacitated edges into a tree (i.e. between every pair of nodes, there is only connecting pathway), while preserving the minimum cuts between all pairs of nodes. Hence, only *k* – 1 cuts need to computed. The effective barrier between minima *i* and *j* is given by: *F_ij_* = – *k_B_T* ln(*N_j_*), where *N_j_* is the number of minimum-cuts between pair *i* and *j*.

**Figure S3.**
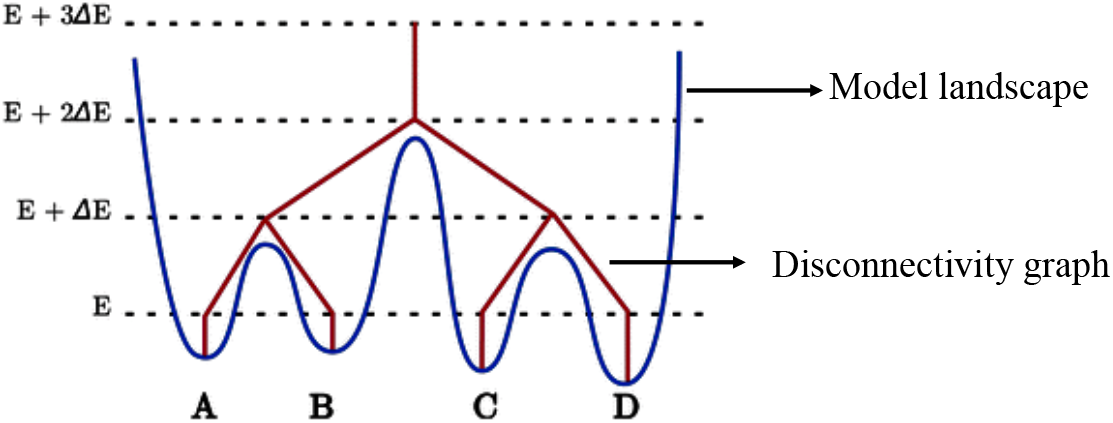
The disconnectivity graph representation (red) of a model free energy landscape (blue). Each local minimum is denoted by a vertical line starting at the energy of that minimum. At any given energy threshold, *E* + ΔE, minima that have an effective free energy barrier below the threshold are grouped into disjoint sets. The figure is adapted from ref.^11^

### Visualizing the free energy landscapes

The free energy landscapes were visualized in the form of transition disconnectivity graphs (TRDGs)^12,13^ In this representation, the landscape is partitioned in disjoint basins of attraction, by choosing an appropriate energy scale (Fig. S2). The local minima within each basin are mutually accessible, whereas minima in different basins are connected by larger free energy barriers. In contrast to other schemes, which rely on low-dimensional projections of the energy landscape onto predefined order parameters, TRDGs preserve the kinetic information. ^7^

In a TRDG, the energy is represented on the vertical axis, while the horizontal axis can be completely arbitrary.^12^ A vertical line is drawn at each minimum (in this case, A-D), beginning at energy of that state. At a threshold of *E* + Δ*E*, minima A and B (and also C and D) are connected together since the effective barriers separating these states are below the threshold. All the minima are connected at level *E* + Δ*E* because the barrier separating the two sets (A and B from C and D) lies below the threshold. The TRDGs were constructed using the disconnectionDPS code. ^14^

### Estimation of landscape roughness density

The ruggedness of the energy landscapes was quantified in terms of roughness density, *ρ_LB_*. It is defined as the number of minima in the TRDG that branch off at a particular energy level.^15^

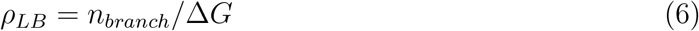

In Eq. 7, *n_branch_* is the percentage of free energy minima that branch off at a particular energy level, and Δ*G* represents the threshold used in the superbasin analysis.

### Definition of N* states

Aggregation-prone conformations (the putative N* state) were identified from the conformational ensembles based on structural overlap with the monomer unit of the experimentally determined A*β*40 and A*β*42 fibril structures. We computed *χ_fib_*(*t*) using:

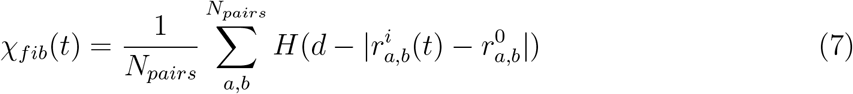

where 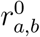 is the distance between sites *a* and *b* in the reference structure. Conformations which exhibit *χ_fib_*(*t*) ≥ *χ_c_* are deemed to be aggregation-prone, and collectively defined as belonging to the N* state. For A*β*40, we chose the monomer unit from the solid-state striated fibril structure (PDB ID: 2M4J) reported by Tycko and coworkers^16^ as the reference state. To identify aggregation-prone conformations in the A*β*42 ensemble, we used the monomer units from two topologically different fibril structures reported by Riek and coworkers using solution-NMR as references, the U-bend fibril structure,^17^ and a S-bend fibril structure.^18^

### Generating all-atom structures from coarse-grained snapshots

The all-atom reconstruction of the coarse-grained (CG) structures was carried out in three stages. (1) First the BBQ code^19^ was used to generate the all-atom representation of the backbone starting from the trace of the C_α_ positions available from the CG structure. (2) Following the backbone reconstruction, the side-chains for each CG conformation was generated using the SCWRL4 formalism.^20^ (3) Finally, the geometry of the reconstructed all-atom structure was further optimized to remove any unwanted steric clashes using the AMBERff12SB force field, in conjunction with the Generalized Born solvent model parametrized by Onufriev, Bashford and Case (GB-OBC).^21^ During the geometry optimization, positional restraints were applied to the *C_α_* atoms to prevent any substantial distortion from the corresponding CG topology. The optimizations were carried out using the *sander* module available within the AMBER12 distribution. ^22^

**Figure S4.**
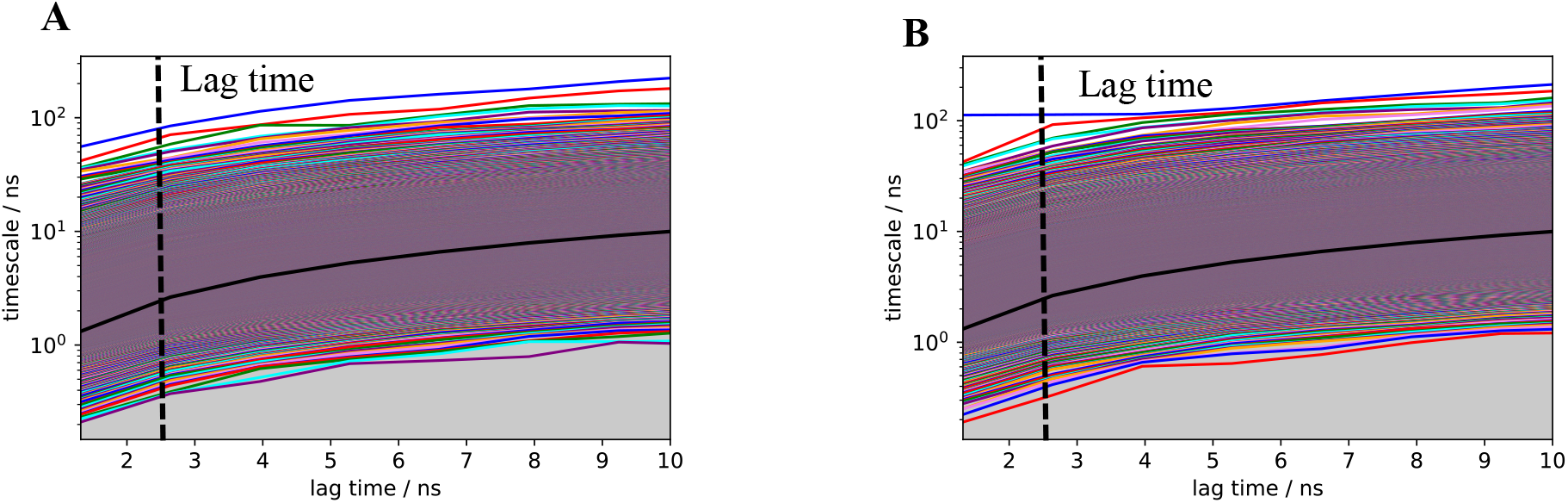
The implied time-scales for A*β*40 (A) and A*β*42 (right) are independent of the lagtime. To compute the relaxation time-scales within the A*β*40 and A*β*42 monomer ensembles, we chose a lagtime of 2.7ns.

**Figure S5.**
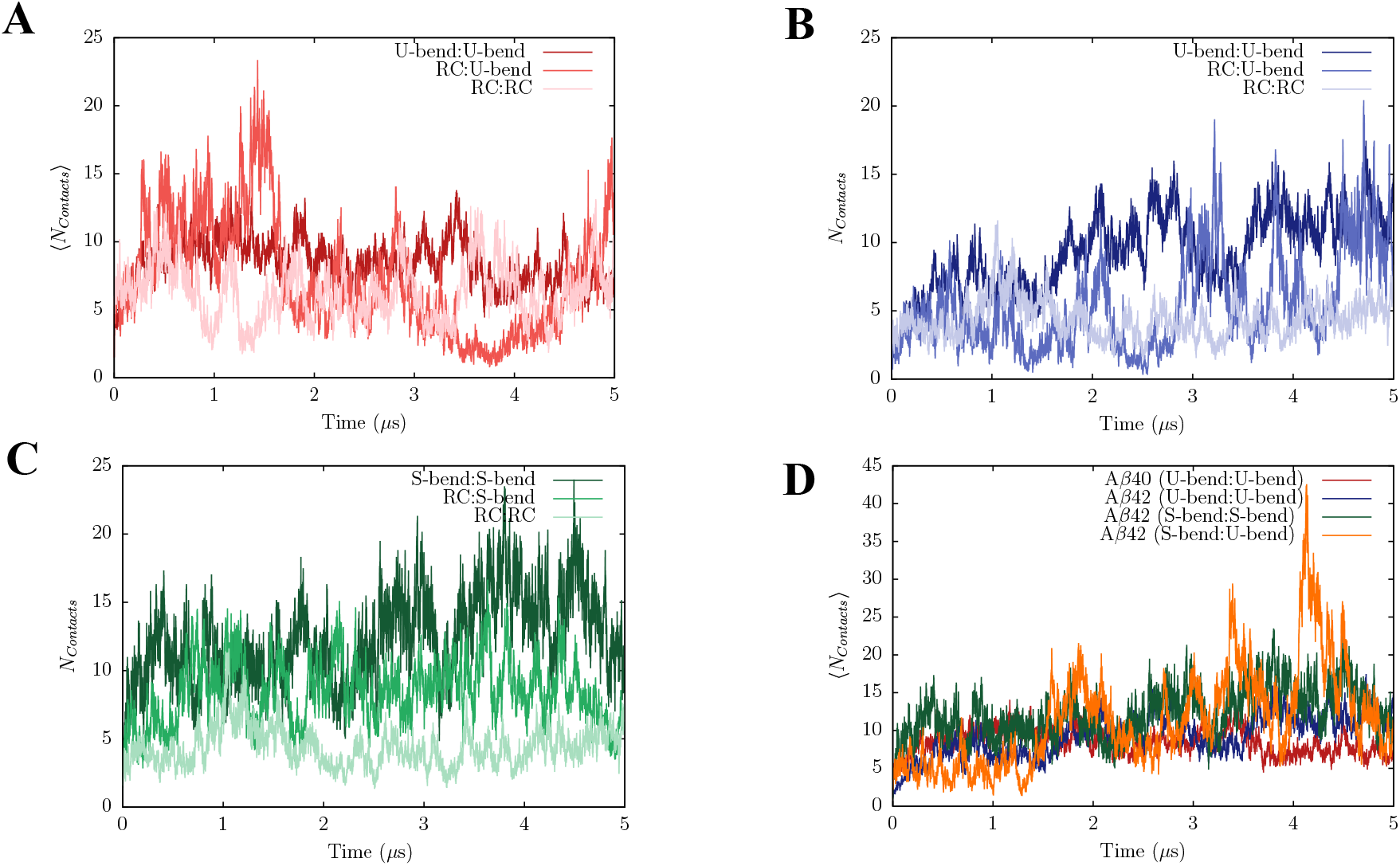
Dimerization profiles of A*β*40 and A*β*42 initiated from different configurations. A dimer is formed when the number of inter-chain contacts, 〈*N_contacts_*〉 exceeds 5. When dimerization reactions are initiated from two RC configurations, there is little or no dimer formation. (B) Just as in A*β*40, dimers formed by a combination of a U-bend structure, and RC configuration, tend to be more dynamic (large fluctuations in 〈*N_contacts_*〉. The propensity to form dimers is rather low, when dimerization reactions are initiated from two RC configurations. (C) The S-bend motif acts as an optimal template for dimer formation (high values of 〈*N_contacts_*〉 throughout the simulation time). Dimers formed by a combination of S-bend and RC-like configurations are also thermodynamically stable compared to those formed by U-bend counterpart. The relatively low values of 〈*N_contacts_*〉 suggest that the tendency to form dimers is rather low when the dimerization is initiated from two RC configurations. (D) Mixed dimers formed by a U-bend and a S-bend motif of A*β*42 are highly dynamic, and can transiently form a large number of interchain contacts.

**Figure S6.**
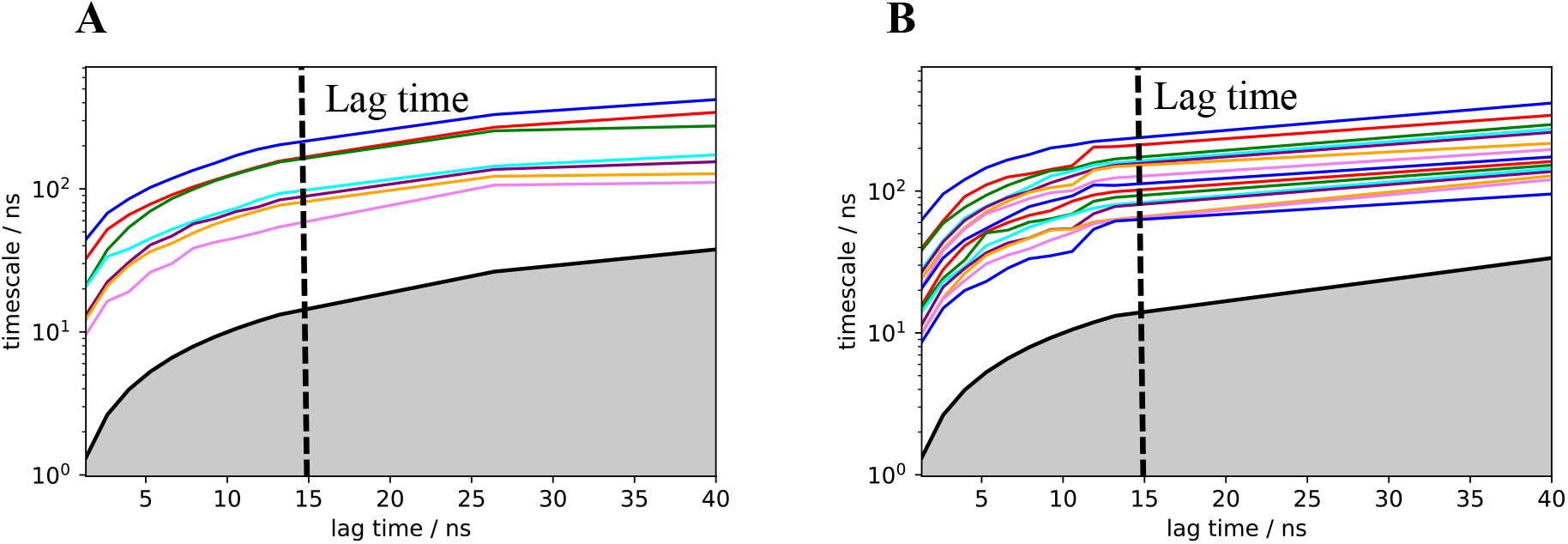
The implied timescales associated with the HMMs for A*β*40 (A) and A*β*42 (B).The transition networks describing the dimerization process were constructed at a lag-time of 15 ns where the timescales appear converged.

